# Brain-resident myeloid cells promote rapid leukocyte adhesion in leptomeningeal vessels after anti-Aβ immunotherapy

**DOI:** 10.64898/2026.09.07.749938

**Authors:** Moustafa Algamal, Shinya Yokomizo, Wadzanai Ndambakuwa, Sarena Abdallah, Maria V. Sanchez-Mico, Rebecca Gillani, Patrick da Silva, Aryan Rajput, Lakshita Palanivelu, Yuna Lee, Katy R. Walsh, Sunny Kumar, Kivilcim Kilic, Joseph El Khoury, Saef Izzy, Mariel G. Kozberg, Brian J. Bacskai

**Author notes:** These authors jointly supervised this work: Brian J. Bacskai, Mariel G. Kozberg. Correspondence should be addressed to Moustafa Algamal.

## Abstract

Anti-amyloid-β (Aβ) immunotherapy improves cognitive outcomes in Alzheimer’s disease (AD) but is associated with amyloid-related imaging abnormalities (ARIA), through poorly understood mechanisms. To define how anti-Aβ antibodies acutely engage brain immune and vascular compartments, we developed a longitudinal in vivo two-photon imaging platform to track microglial dynamics, peripheral immune cell recruitment, and vascular responses in APP/PS1 mice. Anti-Aβ antibodies, including aducanumab and lecanemab, rapidly initiated microglial activation and spatial reorganization within 24 hours of dosing, with recruitment to plaque-associated regions, stabilizing plaque growth. Aducanumab and lecanemab also triggered a rapid and transient cerebrovascular immune response characterized by rolling and adhesion of peripheral immune cells along leptomeningeal vessels, accompanied by endothelial activation. Immune cell characterization revealed recruitment of innate immune cells (Iba1⁺Ki67⁺ monocytes and Ly6G⁺ neutrophils) and proliferative CD3⁺ T cells into the vascular compartment following treatment. Prophylactic treatment with high-dose dexamethasone reduced the number of adherent cells without affecting microglial activation. Depletion of brain-resident immune cells similarly reduced peripheral immune cell recruitment, supporting their contribution to leukocyte recruitment. Postmortem brain tissue from an AD patient treated with lecanemab showed higher proliferation-associated monocyte signature scores, suggesting translational relevance to the proliferative myeloid response observed in our mouse models. These findings demonstrate that anti-Aβ immunotherapy rapidly initiates a coordinated central and peripheral immune response at the leptomeningeal interface. These early immune–vascular interactions represent a plausible initiating mechanism for ARIA and provide a mechanistic framework to guide strategies for mitigating ARIA risk.

## Introduction

Amyloid-β (Aβ) immunotherapies represent a major advance in Alzheimer’s disease (AD), with anti-Aβ monoclonal antibodies demonstrating the ability to reduce cerebral amyloid burden and slow cognitive decline. However, a subset of treated patients develops amyloid-related imaging abnormalities (ARIA), including cerebral edema (ARIA-E) and hemorrhages (ARIA-H), which restrict patient eligibility for treatment and pose significant safety concerns^1^. An incomplete understanding of the mechanism of action of anti-Aβ antibodies in AD, particularly the cellular and molecular mechanisms linking amyloid clearance to vascular and innate immune responses, is a barrier to developing therapeutic strategies to prevent and treat ARIA.

Both central and peripheral immune responses may critically influence the therapeutic efficacy and vascular side effects of anti-Aβ immunotherapy^2–5^. Postmortem human studies of both active and passive Aβ immunization highlight the central role of microglia in mediating Aβ clearance^2^, while growing evidence implicates the recruitment of peripheral immune cells in ARIA pathogenesis. Postmortem analysis of a patient who died from cerebral amyloid angiopathy-related inflammation following lecanemab treatment revealed perivascular lymphocytic infiltrates, CD68⁺ macrophage accumulation, and fibrinoid degeneration of the vessel walls^3^. Single-cell profiling of peripheral blood from ARIA⁺ patients identified a selective expansion of cytotoxic CD8⁺ T cells together with enhanced antigen-presentation signaling by CD14⁺ and CD16⁺ monocytes, suggesting increased monocyte–T cell communication in ARIA^+^ patients^5^. Animal studies further support a role of peripheral immune recruitment in anti-Aβ antibody-mediated vascular responses. Mice receiving chronic anti-Aβ antibody (3D6) treatment exhibit colocalization of monocytes and perivascular macrophages at sites of microhemorrhage^6^. Additionally, TREM2⁺ monocyte-derived perivascular macrophages, CD4 and CD8 cells accumulate at sites of vascular amyloid deposition following antibody treatment^7^.

Despite growing evidence implicating both central and peripheral immune populations in anti-Aβ antibody-mediated amyloid clearance and vascular injury, the temporal sequence of events linking antibody administration to immune activation, vascular inflammation, and tissue injury remains poorly defined. It is unknown which immune populations respond first, how central and peripheral immune responses are coordinated, and how these early events contribute to subsequent amyloid clearance and vascular pathology. Defining this sequence and the type of immune cells involved at each stage is critical for optimizing treatment strategies that maximize plaque clearance while minimizing vascular side effects.

We employed longitudinal two-photon imaging, flow cytometry and immunohistochemistry in APP/PS1 mice to dissect central and peripheral immune cell dynamics in the brain and neurovascular interface following anti-Aβ antibody administration. We found that microglia migrate toward plaques and proliferate within 24 hours of treatment. Additionally, we observed rapid and robust acute vascular inflammation, characterized by the recruitment of monocytes, neutrophils and T-cells within hours of antibody administration. A significant proportion of monocytes and T-cells were proliferative and were detected in brain parenchyma. An exploratory human transcriptomic analysis provided evidence consistent with a proliferation-associated myeloid response in postmortem human brain tissue from a patient who developed neurological symptoms shortly after receiving lecanemab and died after treatment with tPA for presumed ischemic stroke and subsequently was found to have vascular inflammation by postmortem analysis. Together, these findings indicate that while anti-Aβ antibodies effectively engage microglia, promoting plaque clearance, but trigger peripheral immune cell recruitment that may compromise vascular integrity. This work provides mechanistic insight into anti-Aβ immunotherapy-associated adverse events and establishes an in vivo platform for testing strategies to mitigate ARIA risk.

## Results

### Aducanumab administration leads to rapid microglial recruitment to Aβ plaques and prevents plaque growth in APP/PS1 mice

To determine how anti-Aβ immunotherapy influences microglial behavior, we performed longitudinal in vivo two-photon imaging through chronic cranial windows in APP/PS1 mice expressing a Tmem119-tdTomato microglial reporter. APP/PS1 mice were crossed with Tmem119-tdTomato-CreERT2 mice, enabling specific visualization of resident microglia based on TMEM119 expression, which is absent in infiltrating peripheral macrophages^8^. Validation of reporter specificity confirmed selective labeling of microglia (Extended Data Fig. 1).

Amyloid plaque was visualized by the injection of methoxy-X04 24 hours (h) prior to each imaging session. Baseline microglial spatial organization was established prior to treatment, followed by repeated imaging after systemic administration of either IgG control or aducanumab (Fig. 1a). Weekly imaging revealed stable microglial in IgG-treated mice, with most microglia maintaining consistent positions across imaging sessions (Fig. 1b). In contrast, aducanumab treatment was associated with pronounced microglial spatial rearrangements, including loss of previously identified microglia from the imaging field and the appearance of new microglia at distinct locations.

**Figure 1.**
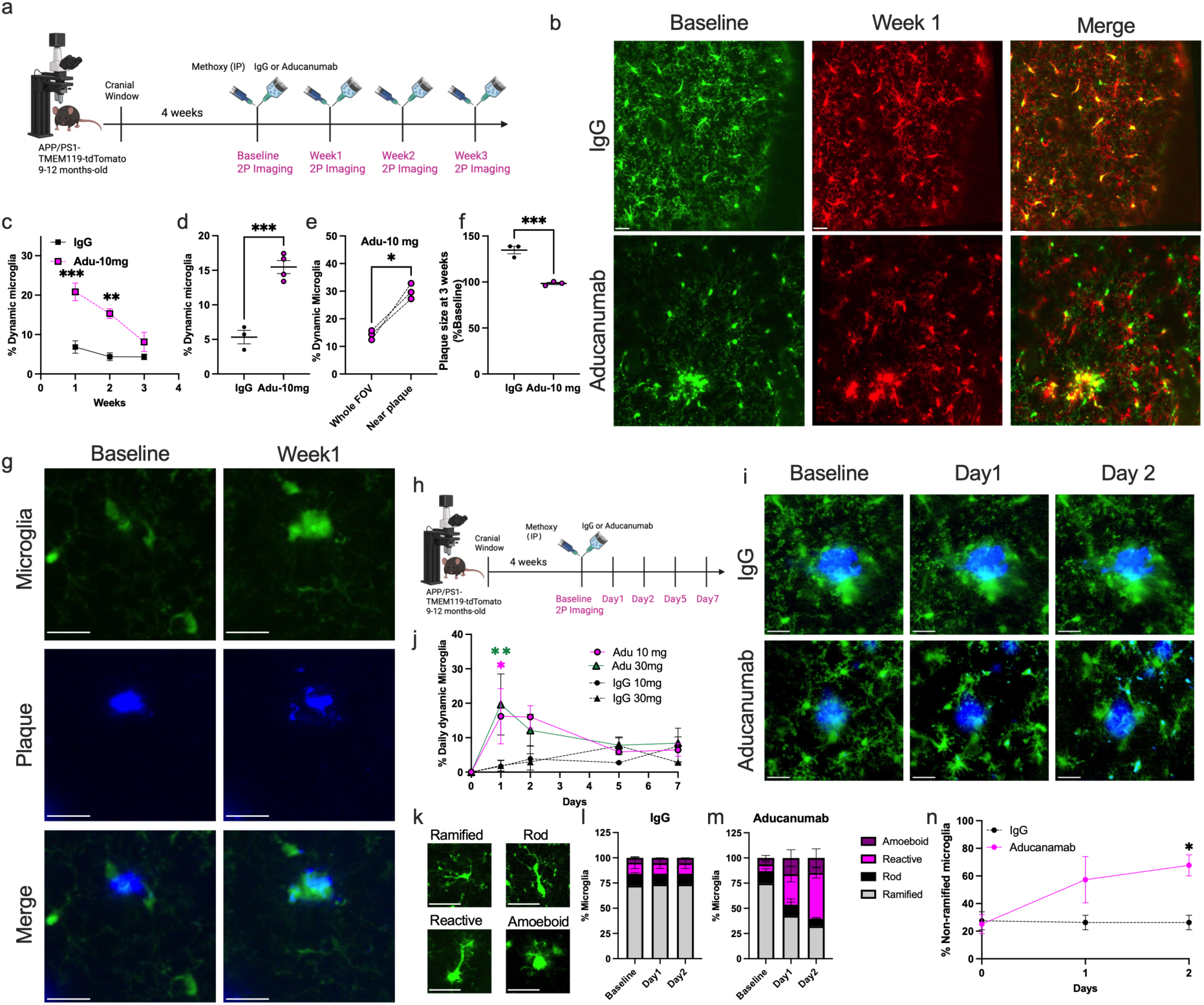
Aducanumab promotes early microglial recruitment and suppresses amyloid plaque growth. (a) Experimental design: APP/PS1; Tmem119-tdTomato mice underwent cranial window implantation, baseline imaging, systemic IgG or aducanumab (10 mg/kg) and weekly two-photon imaging for 3 weeks. (b) Representative images of microglia at baseline and during follow-up imaging sessions after IgG or aducanumab injection. (c) Weekly changes in microglial dynamics, expressed as the average percentage of lost and newly appearing microglia per imaging session. Repeated-measures mixed-effects model (REML), P < 0.0001; n = 3-4 mice per group) (d) Average weekly microglial dynamics measured over 3 weeks of longitudinal imaging (unpaired two-sided t-test, P=0.0007, n = 3-4 mice per group). (e) Increased microglial dynamics within 50 μm of plaques relative to the whole field of view following treatment (Paired two-sided t-test, P=0.022; n =3 mice). (f) Plaque size normalized to baseline at 3 weeks post-treatment with 10 mg/Kg aducanumab (unpaired two-sided t-test, P=0.001, n = 3 mice per group). (g) Representative in vivo images illustrating amyloid plaque morphology following aducanumab administration; microglia (green) and amyloid plaques (blue) are shown at baseline and at one week post-treatment. (h) Schematic of a daily multiphoton imaging paradigm in APP/PS1; Tmem119-tdTomato mice, with cranial window implantation, baseline imaging, IgG or aducanumab (10 or 30 mg/kg), and 7 days of imaging. (i) Representative images (baseline, Day 1, and Day 2) showing increased plaque-associated microglial dynamics after aducanumab. (j) Quantification of daily dynamic microglia over 7 days (REML, Main time effect P=0.007, n= 3 mice per group). (k) Representative examples of microglial morphology classification. Microglia were categorized as ramified, rod-shaped, reactive, or amoeboid^10^. (l) Minimal changes in microglial morphology were observed following IgG injection, whereas the proportion of amoeboid and reactive microglia increased following antibody administration (m). (n) Increased percentage of non-ramified microglia, defined as the combined proportion of rod-shaped, reactive, and amoeboid microglia, within 1–2 days following aducanumab treatment (two-way ANOVA, main effect of treatment, *P* = 0.026; *n* = 2-3 mice per group). Two-way ANOVA was followed by Holm–Šídák’s multiple-comparisons test. Scale bars, 20 µm. \**P* < 0.05, \*\**P* < 0.01, \*\*\**P* < 0.001. Schematics in (a) and (h) were created with BioRender.com.

Quantification across animals demonstrated a significant increase in the proportion of microglia that were either lost from their original location or newly appeared over a 3-week period in aducanumab-treated mice compared to IgG controls (Extended Data Fig. 2a, b). The average proportion of newly appearing and lost microglia was defined as “dynamic microglia” and used throughout the manuscript as a measure of microglial reactivity.

When analyzed on a per-week basis, aducanumab treatment induced a marked increase in microglial dynamics potentially due to repositioning of microglia, although our methods are unable to distinguish between a repositioned or newly formed microglia (Fig. 1c). Average weekly microglial dynamics was significantly elevated in aducanumab-treated mice over 3 weeks, whereas IgG-treated mice showed minimal long-term change (Fig. 1d). Switching IgG-treated mice to aducanumab induced a significant increase in microglial dynamics (Extended Data Figure 2), indicating that baseline amyloid plaque burden did not account for the observed effects. These findings suggest that anti-Aβ immunotherapy induces sustained alterations in microglial dynamics. Consistent with the known effects of aducanumab on amyloid pathology^9^, microglial dynamics were particularly pronounced in plaque-proximal regions, with significantly increased dynamic microglia near plaques (Fig. 1e, Movies 1-3). Quantitative analysis confirmed that plaque size remained stable over the 3-week aducanumab treatment period, whereas plaques continued to grow in IgG-treated mice over the same imaging interval (Fig. 1f, g). To further resolve the temporal dynamics of microglial rearrangement, we performed a separate experiment using daily rather than weekly imaging following treatment (Fig. 1h). Microglial dynamics changed within 24– 48 h after antibody administration (Fig. 1i, j). This was accompanied by a shift toward non-ramified microglial morphologies, including amoeboid and reactive states (Fig. 1k-n). Together, these data demonstrate that aducanumab treatment induces robust microglial recruitment and activation, particularly in plaque-associated regions, and that these changes occur within 24 h of antibody administration.

### Aducanumab induces rapid and transient intravascular peripheral immune cell recruitment

While performing high–temporal resolution two-photon imaging to track microglial dynamics, with blood vessels labeled by intravenous dextran-fluorescein, we unexpectedly observed numerous circulating cells exhibiting rolling and firm adhesion along meningeal veins 18 h after aducanumab administration (Fig. 2a; Movie 4). These cells were identified by their exclusion of 70 kDa dextran (Movie 4). This intravascular behavior was absent at baseline and in IgG-treated controls (Fig. 2a). Quantification across animals showed a significant increase in rolling or adhering intravascular peripheral immune cells at 18 h following aducanumab compared with IgG treatment (Fig. 2b). Notably, the appearance of these cells was temporally coincident with the early microglial recruitment, prompting us to examine whether anti-Aβ immunotherapy acutely engages cerebrovascular immune responses. To identify rolling and adherent intravascular cells, mice received intravenous dextran-fluorescein prior to baseline imaging to label the blood plasma. The same vascular field of view was imaged at baseline and again 18 h after aducanumab administration. Following the 18 h imaging session, Lycopersicon esculentum (LE) lectin was injected and imaged in vivo, after which animals were immediately euthanized and perfused, and the same vessels were examined by whole-mount ex vivo staining (Fig. 2c). Although LEL is commonly used to label endothelial cells, it can also bind to leukocytes^11^. Low- and high-magnification in vivo imaging confirmed firm leukocyte adhesion to the vessel wall (Fig. 2d, e, g). Corresponding ex vivo lectin staining verified the intravascular localization of these cells (Fig. 2f, g). Interestingly, these cells were frequently observed near perivascular macrophages within meningeal veins and were also detected outside the vasculature in cerebral amyloid angiopathy (CAA)-affected vessels (Extended Data Figure 3).

**Figure 2.**
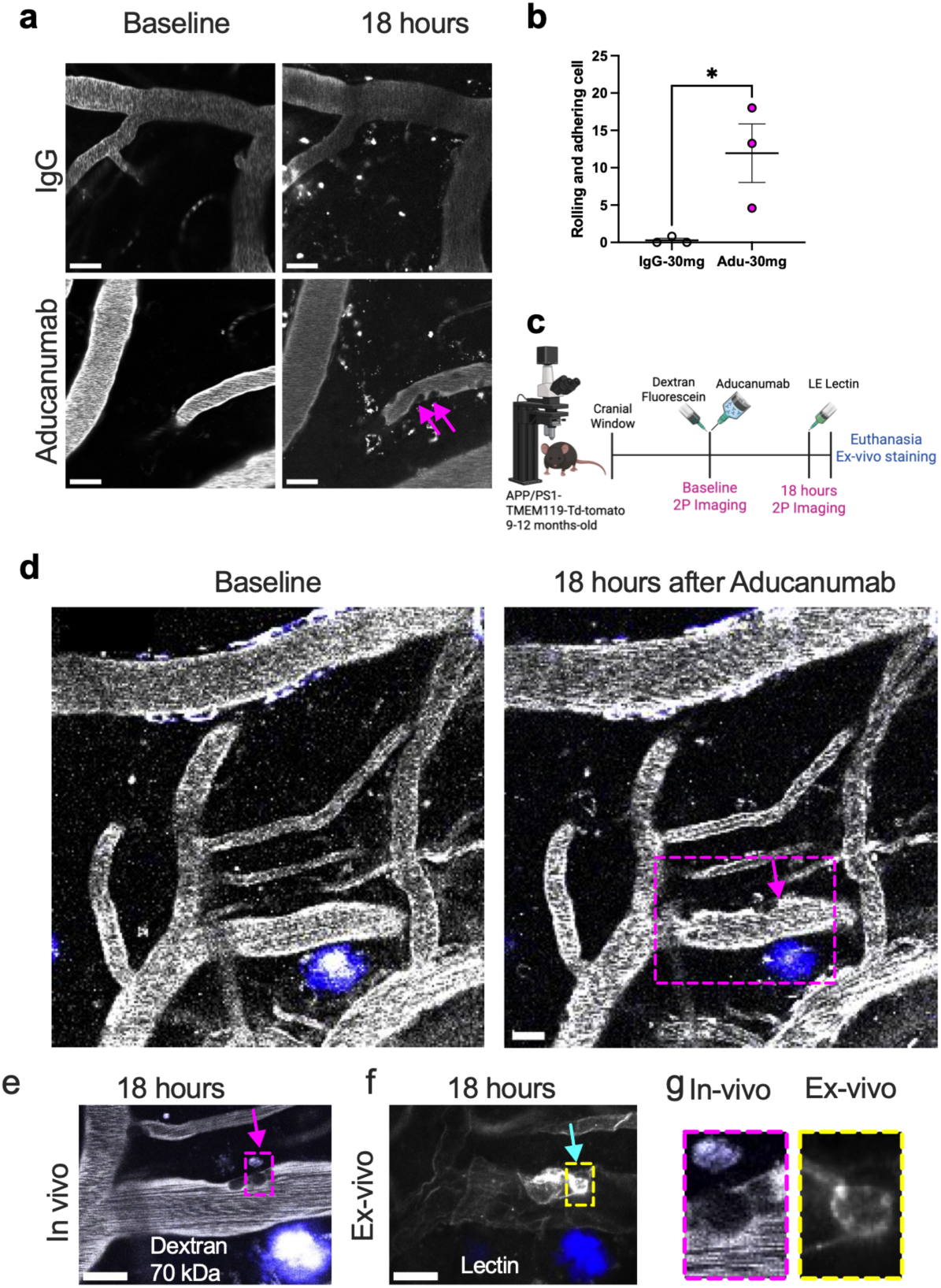
Anti-Aβ antibodies induce rapid intravascular leukocyte rolling and adhesion in leptomeningeal vessels. **(a)** Representative in vivo two-photon images of meningeal venous vessels at baseline and 18 h after administration of IgG or aducanumab. Arrowheads indicate intravascular cells exhibiting rolling or firm adhesion that emerge following aducanumab treatment. **(b)** Quantification of intravascular peripheral immune cells at 18 h demonstrates a significant increase following aducanumab compared with IgG controls (unpaired two-sided t-test, P=0.041, n = 3 mice per group). **(c)** Experimental timeline for acute vascular imaging following aducanumab treatment. APP/PS1 mice (9–12 months old) implanted with chronic cranial windows received intravenous dextran-fluorescein prior to baseline imaging to label blood vessels, followed by aducanumab administration, repeat two-photon imaging at 18 h, and subsequent euthanasia for ex vivo lectin staining. **(d)** Low-magnification in vivo images of the same meningeal vascular network at baseline and 18 h after aducanumab administration. Dashed box denotes the region shown at higher magnification. **(e)** High-magnification in vivo image illustrating an adherent intravascular leukocyte (arrow) along the venous vessel wall after aducanumab treatment. **(f)** Corresponding ex vivo lectin staining confirms the intravascular localization of the adherent leukocyte (arrow) observed in vivo. **(g)** Side-by-side comparison of the same cell imaged in vivo and ex vivo. Scale bars, 20 µm. Schematic in (c) was created with BioRender.com.

### Lecanemab administration leads to rapid microglial and peripheral immune cell recruitment in APP/PS1 mice

Given that lecanemab is an anti-Aβ antibody now in clinical use and has a distinct epitope preference and clinical profile compared to aducanumab, we next asked whether the rapid alteration in microglial spatial rearrangements represents a general response to anti-Aβ immunotherapy rather than an antibody-specific effect. To address this, we performed longitudinal in vivo two-photon imaging after lecanemab administration. Tmem119-tdTomato; APP/PS1 mice were imaged at baseline and then at 7, 18 and 48 h post-treatment to track microglia and lectin⁺ cells (Fig. 3a). To monitor peripheral immune cell dynamics, intravenous LE lectin was administered prior to each imaging session. Lecanemab administration rapidly increased microglial dynamics, peaking at 18 h and declining by 48 hours (Fig. 3b-e). In parallel, lecanemab administration induced robust intravascular leukocyte rolling and adhesion within meningeal venous vessels, which were detectable as early as 7 h after treatment and declined by 48 h (Fig. 3c, f-g, Movie 5). Quantitative analysis revealed a rapid and transient increase in both rolling and adherent lectin-positive leukocytes (Fig. 3f, g) following lecanemab, while IgG-treated mice exhibited minimal changes across all time points (Fig. 3e–g). Consistent with this, switching from IgG to lecanemab increased intravascular rolling cells within the same vessels over time (Extended Data Figure 4). Together, these data demonstrate that anti-Aβ antibodies rapidly and transiently induce intravascular leukocyte recruitment and adhesion within meningeal vessels. To determine whether lecanemab alters microglial activation states, APP/PS1 mice (10–13 months old) received a single injection of lecanemab or IgG and were analyzed 18 h later by immunofluorescence (Fig. 3h). Lecanemab-treated mice showed reduced microglial branch length (Fig. 3j) and ramification index (Fig. 3k), consistent with rapid morphological activation, together with an increased number of parenchymal IBA1⁺Ki67⁺ cells, with Ki67 serving as a marker of cell proliferation (Fig. 3l). These findings indicate that microglia rapidly adopt an activated and proliferative phenotype following anti-Aβ immunotherapy.

**Figure 3.**
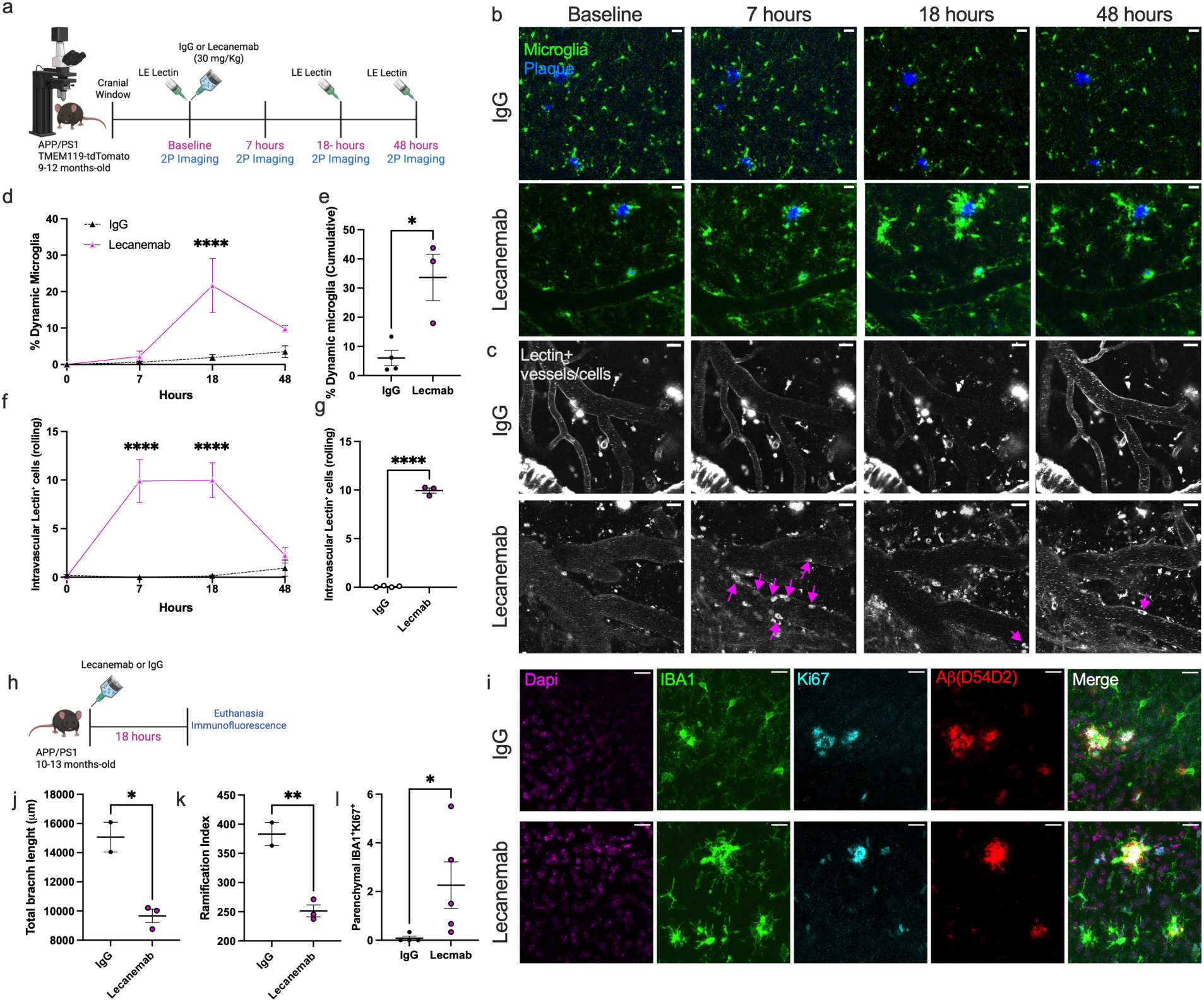
Daily in vivo imaging reveals rapid microglial dynamics following lecanemab treatment. **(a)** Experimental design schematic. Tmem119-tdTomato; APP/PS1 mice implanted with chronic cranial windows underwent baseline two-photon (2P) imaging of cortical microglia, followed by systemic administration of 30 mg/Kg lecanemab and imaging at 7-, 18-, and 48-hours (h) post-treatment. LE lectin was used to visualize peripheral immune cells **(b)** Representative in vivo 2P images illustrating microglial organization at baseline and during daily follow-up imaging after lecanemab treatment. (c) Time course of intravascular lectin⁺ rolling and adhering leukocytes show a rapid increase after lecanemab, peaking at 7 h and declining by 48 h, with minimal change in IgG controls. Arrowheads indicate rolling and adherent intravascular leukocytes observed after lecanemab treatment. (d) Quantification of microglial dynamics at each time point following lecanemab administration (Two-way repeated-measures ANOVA, main treatment effect: P = 0.013; n = 3–4 mice per group) **(e**) Cumulative microglial dynamics through Day 2, expressed as the sum of interval-specific percentages of dynamic microglia (unpaired two-sided t-test, P = 0.014; n = 3–4 mice per group). **(f)** Quantification of intravascular lectin-positive rolling and adhering cells at different time points confirms increased leukocyte rolling and adhesion in lecanemab-treated mice relative to IgG controls (Two-way repeated-measures ANOVA, main treatment effect: P <0.001; n = 3–4 mice per group). **(g)** Average number of rolling and adhering cells per mouse after 18 h of antibody administration (unpaired two-sided t-test, P<0.0001, n = 3-4 mice per group). **(f)** Experimental design: APP/PS1 mice (10–13 months) received lecanemab or IgG and were analyzed 18 h later by immunofluorescence **(i)** Representative immunofluorescence images from IgG- and lecanemab-treated APP/PS1 mice stained for DAPI (magenta), IBA1 (green), Ki67 (cyan), and amyloid plaques (red). Lecanemab treatment induced clustering of activated plaque-associated microglia with increased Ki67 immunoreactivity adjacent to amyloid plaques. Quantification of microglial total branch length **(j)**, microglial ramification index **(k)** and the number of parenchymal IBA1^+^Ki67^+^ cells **(l)** following treatment (Unpaired two-sided *t*-test, *P* = 0.01, 0.007, and 0.022, respectively; *n* = 2–5 mice per group). Data are shown as mean ± SEM. Two-way repeated-measures ANOVA was followed by Holm–Šídák’s multiple-comparisons test. Statistical significance is indicated (*p<0.05, **p<0.01, ***p<0.001, ****p<0.0001). Scale bars, 20 µm. Schematics in (a) and (h) were created with BioRender.com.

### Lecanemab induces rapid leptomeningeal adhesion of monocytes, neutrophils and T cells in APP/PS1 mice

To define the identity of peripheral immune cells recruited after anti-Aβ immunotherapy to the cerebral vasculature, APP/PS1 mice (10–13 months old) received a single injection of lecanemab or IgG control and were analyzed 18 h later by immunofluorescence (Fig. 4a). Staining for anti-Aβ and anti-IgG confirmed vascular amyloid and plaque-associated localization of lecanemab that was absent in IgG-treated controls (Fig. 4b). We first assessed recruitment of CD43+ cells, an adhesion marker expressed on several leukocytes in leptomeningeal and cortical vessels. Lecanemab-treated mice showed a striking increase in meningeal CD43+ cells compared with IgG controls, with similar accumulation observed along cortical vessels (Fig. 4c). CD43⁺ cells were observed in both veins and SMA⁺ arteries (Fig. 4e). Lecanemab treatment also increased the number of parenchymal CD43⁺Ki67⁺ cells (Fig. 4f), consistent with the presence of proliferation-marker-positive leukocytes outside the vascular compartment. Together, these findings support rapid leukocyte accumulation at the cerebrovascular interface following antibody administration. To further characterize the identity and origin of these cells, we stained for Iba1, a marker predominantly expressed by myeloid cells, including peripheral monocytes, border macrophages, and brain microglia. Lecanemab treatment increased the number of Iba1⁺Ki67⁺ cells in leptomeningeal vessels compared with controls (Fig. 4g–i), suggesting expansion and/or recruitment of proliferative myeloid cells in perivascular niches following anti-Aβ treatment.

**Figure 4.**
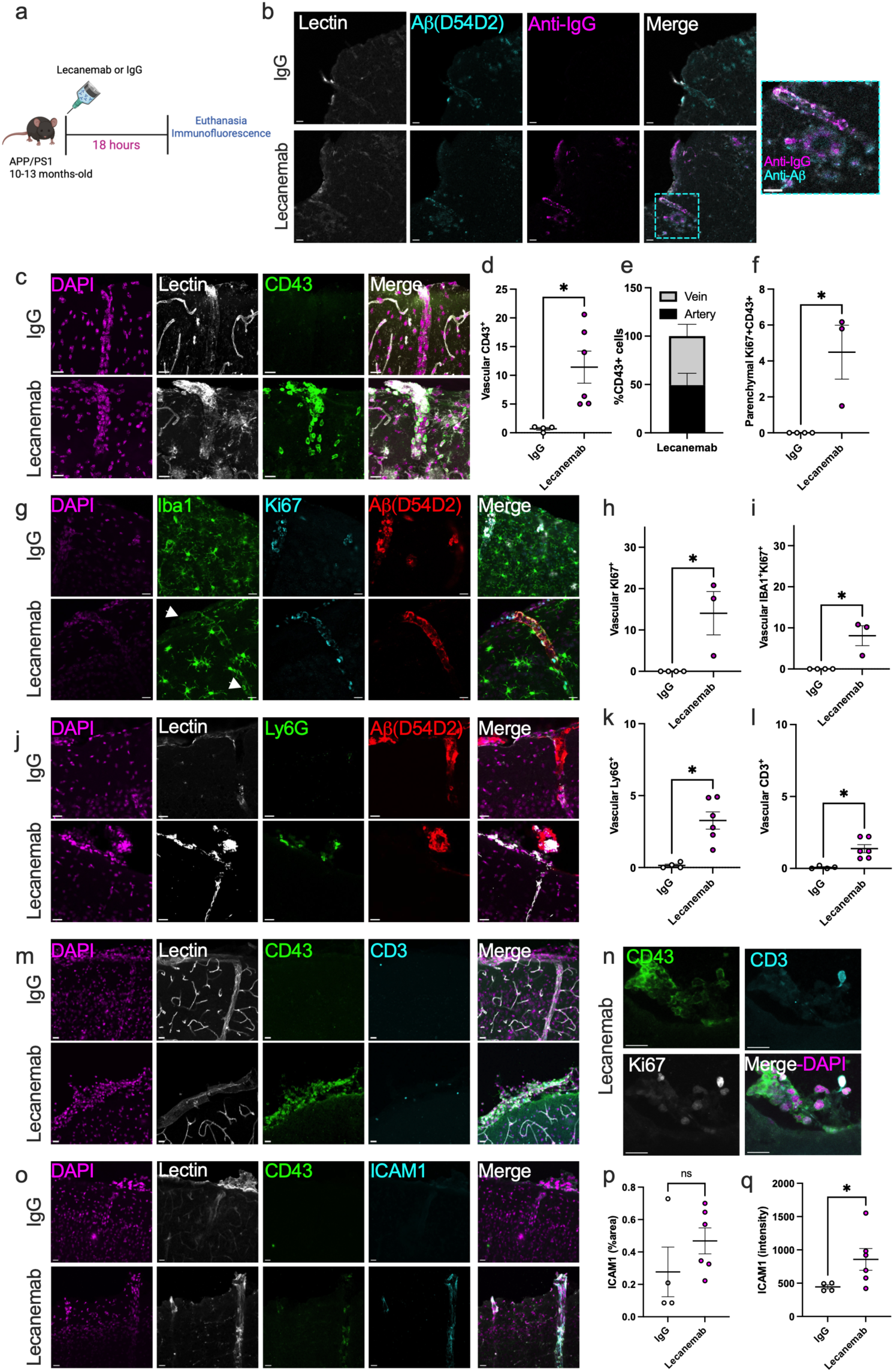
Lecanemab induces rapid vascular and immune activation with recruitment of peripheral immune cells. (a) Experimental design: APP/PS1 mice (10–13 months) received lecanemab (30mg/kg) or IgG and were analyzed 18 h later by immunofluorescence. (b) Representative images showing anti-Aβ and anti-IgG localization with LE lectin labeling, demonstrating antibody engagement with vascular and parenchymal Aβ. (c) Representative images of CD43⁺ immune cells adhering to leptomeningeal vessels following lecanemab treatment. Quantification demonstrated increased vascular-associated CD43⁺ cells (d) after lecanemab administration (unpaired two-sided t-test; adjusted P = 0.045, n = 4–6 mice per group). (e) Relative percentages of adhering CD43⁺ cells in veins and arteries. (f) Increased parenchymal Ki67⁺CD43⁺ cells 18 hours after lecanemab (unpaired two-sided t-test; adjusted P = 0.045, n = 3–4 mice per group). (g) Representative images demonstrating vascular-associated Iba1⁺Ki67⁺ proliferating myeloid cells following lecanemab treatment (arrowheads). (h, i) Quantification of vascular-associated Ki67⁺ cells (h) and IBA1⁺Ki67⁺ myeloid cells (i) following lecanemab or IgG treatment (unpaired two-sided t-tests; P = 0.046 and 0.040, respectively; n = 3–4 mice per group) (j) Representative images of neutrophils (Ly6G⁺) adhering to leptomeningeal vessels following lecanemab treatment. (k) Increased recruitment of Ly6G⁺ cells to vasculature following lecanemab treatment (unpaired two-sided t-test; P = 0.02; n = 4–6 mice per group). (l) Increased accumulation of vascular-associated CD3⁺ T cells following lecanemab administration treatment (unpaired two-sided t-test; P = 0.03; n = 4–6 mice per group). (m) Representative images of CD3⁺ cells adhering to leptomeningeal vessels following lecanemab treatment. (n) Example of CD3^+^ CD43^+^ Ki67^+^ T cell adhering to a leptomeningeal vessel after lecanemab treatment. (o) Representative images of ICAM1 expression in leptomeningeal vessels. (p) Average percentage of ICAM1-positive area in lecanemab- and IgG-treated mice (Mann-Whitney U test, P=0.23). (q) Increased ICAM1 intensities in lecanemab-treated mice compared with IgG-treated mice (Mann-Whitney U test, P=0.038), n= 4-6 mice, suggesting endothelial activation. Data are presented as mean ± SEM with individual animals shown. Scale bars, 20 µm. *P < 0.05. P values for t-tests were corrected for multiple comparisons using the Holm–Šídák method. Schematic in (a) was created with BioRender.com.

We then asked whether granulocytic cells were similarly recruited. Ly6G staining revealed a significant increase in Ly6G+ cells in the meninges and along cortical vessels in lecanemab-treated mice compared with IgG-treated animals (Fig. 4j–k), indicating that neutrophil-like cells are also mobilized to the vascular compartment in response to treatment. In addition to myeloid cells, lecanemab increased the accumulation of CD3⁺ T cells along leptomeningeal vessels compared with controls (Fig. 4l–m). A subset of these vascular-associated T cells expressed the proliferation marker Ki67 (Fig. 4n), consistent with the recruitment of proliferative T cells to sites of vascular immune activation. Finally, to determine whether endothelial activation was associated with peripheral immune cell adhesion, we assessed vascular ICAM1 immunoreactivity. Lecanemab treatment was associated with increased ICAM1 staining intensity relative to IgG controls (Fig. 4o–q), consistent with endothelial activation.

We next examined whether these cerebrovascular immune changes were accompanied by alterations in circulating myeloid cells by performing single-cell RNA sequencing of blood monocytes isolated by negative selection, collected 18 h after lecanemab or IgG administration. Quantification of monocyte subsets revealed a significant increase in non-classical monocytes following lecanemab treatment compared with IgG controls (Extended Data Fig. 5). These findings indicate that anti-Aβ immunotherapy rapidly alters the peripheral monocyte compartment, enriching for circulating non-classical monocytes that may contribute to the vascular immune response. Together, these data demonstrate that a single dose of lecanemab rapidly induces cerebrovascular immune activation characterized by endothelial activation and recruitment of multiple peripheral immune cell populations, including CD43+ cells, Ly6G+ granulocytes, Iba1+Ki67+ myeloid cells, and CD3+ T cells. These cerebrovascular changes are accompanied by a shift in peripheral monocytes toward a non-classical phenotype. The Iba1+Ki67+ myeloid cells may represent recently proliferated bone marrow-derived monocytes or a distinct transitional (Tr) proliferative monocyte subset, characterized by high expression of cell cycle-associated genes ^12^.

### Higher proliferation-associated monocyte signature scores in a lecanemab-treated postmortem AD case

We next asked whether a similar proliferative Ki67+ monocyte population could be identified in human tissue after lecanemab administration. To address this, we analyzed publicly available single-cell RNA-sequencing data from postmortem brain tissue of a 65-year-old woman with Alzheimer’s disease who developed neurological symptoms four days after receiving lecanemab^2,4^. She was treated with tissue plasminogen activator (tPA) for a presumed ischemic stroke but subsequently developed intracerebral hemorrhages and died several days later. Autopsy revealed severe cerebral amyloid angiopathy-related inflammation (CAA-ri).^4^ Single-cell RNA-seq analysis^2^ of this case, together with three untreated AD controls, identified the major brain cell populations by UMAP visualization (Fig. 5a). Monocytes in the lecanemab-labeled samples showed higher TR-Mo signature scores and increased expression of proliferation-associated genes, including MKI67, TOP2A, CDK1, BIRC5, and UBE2C, compared with untreated AD samples (Fig. 5b–c). Pathway analysis of the TR-Mo-associated population identified enrichment of pathways related to antigen processing and presentation, interferon and cytokine signaling, cell-cycle regulation, and innate immunity (Fig. 5d). Ligand–receptor analysis predicted interactions involving this population and other immune and vascular cell populations, including APOE–TREM2, APP–CD74, and SPP1–CD44 signaling (Fig. 5e). These exploratory findings suggest a proliferation-associated myeloid response in this case, although its relationship to lecanemab treatment requires confirmation in additional patients.

**Figure 5.**
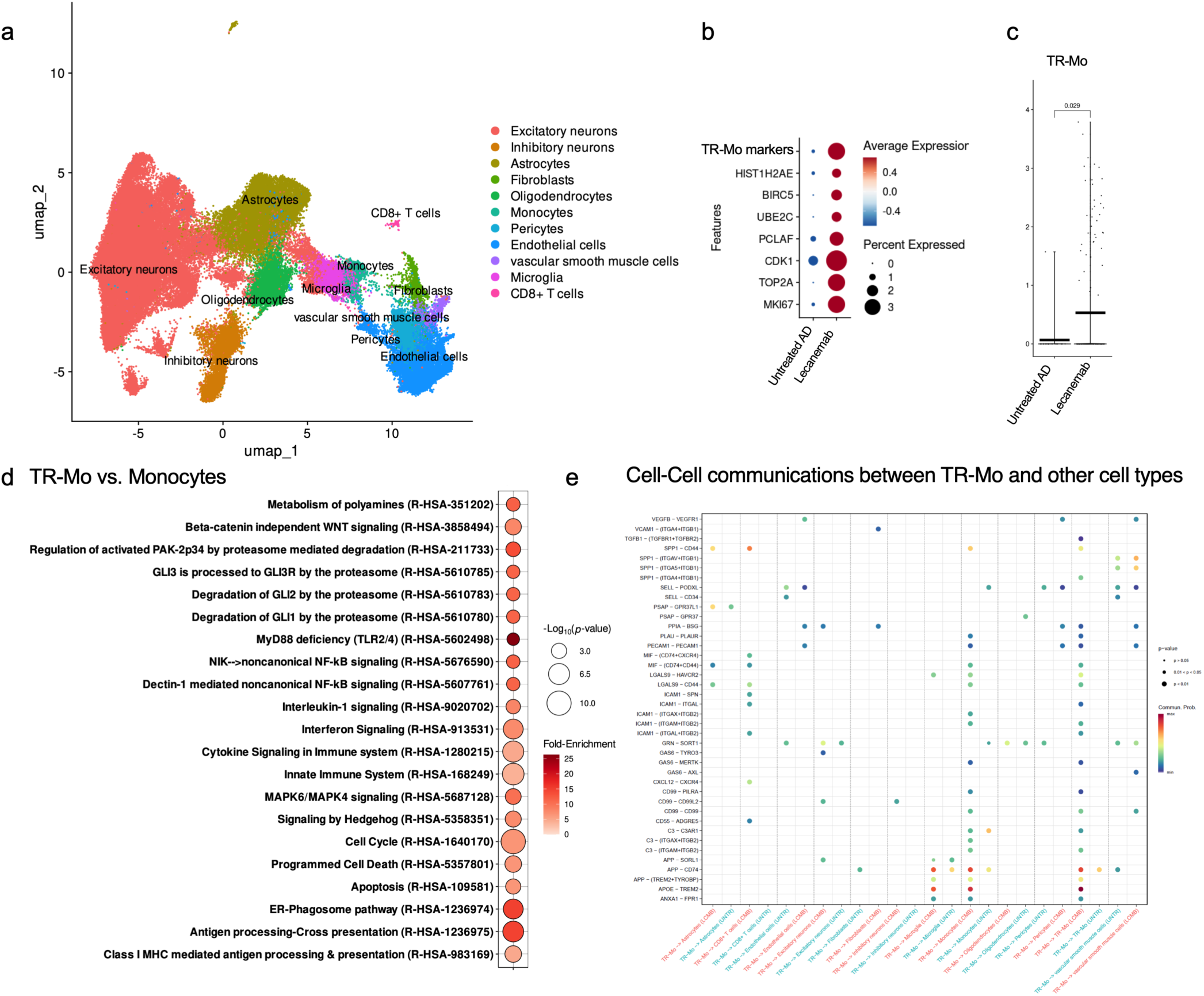
Higher proliferation-associated monocyte signature scores in a lecanemab-treated postmortem AD case. (a) UMAP projection of human brain single-cell RNA-seq data colored by major cell types. (b) Dot plot showing the expression of transitional monocyte (TR-Mo) markers and proliferation-associated genes (HIST1H2AE, BIRC5, UBE2C, PCLAF, CDK1, TOP2A, MKI67) in monocytes from three untreated patients with AD and one lecanemab-treated patient. Dot size represents the percentage of cells expressing each gene, and color represents average scaled expression. (c) Distribution of TR-Mo signature scores within monocytes from lecanemab-labeled and untreated AD samples (d) Pathway enrichment analysis of TR-Mo identifies enrichment of antigen processing and presentation, interferon and cytokine signaling, cell-cycle regulation, and innate immune pathways. (e) Statistically significant cell–cell communication events involving TR-Mo inferred from ligand– receptor analysis, highlighting altered interactions with vascular and immune cell populations in the lecanemab-treated patient.

### Dexamethasone treatment and depletion of brain-resident myeloid cells each attenuate lecanemab-induced peripheral immune cell recruitment

To investigate the mechanisms underlying peripheral immune cell recruitment to the leptomeningeal vasculature, we next asked whether brain-resident myeloid cells and inflammatory signaling contribute to this response. We therefore used two complementary approaches: depletion of brain-resident myeloid cells with PLX5622 and pharmacological suppression of peripheral inflammation with dexamethasone before lecanemab administration. Aged APP/PS1 mice (13–17 months old) were maintained on either a control or PLX5622-containing diet for 6 days. A separate group received dexamethasone (4 mg/kg) every 8 h beginning 1 h before lecanemab (30 mg/kg) administration, and all mice were analyzed 18 h later (Fig. 6a). Flow cytometric analysis of peripheral blood revealed that dexamethasone reduced the proportion of circulating monocytes and increased the proportion of neutrophils, whereas PLX5622 treatment did not significantly alter circulating immune cell populations (Fig. 6b–e). Consistent with efficient depletion of border-associated macrophages (BAMs), PLX5622 markedly reduced CD206^+^ BAM density in the leptomeninges (Fig. 6f, g). PLX5622 treatment also reduced microglial density and total branch length (Fig. 6h–j), but not ramification (Fig. 6k), confirming effective microglial depletion. Dexamethasone had no significant effect on BAM density, microglial density, or microglial morphology (Fig. 6g–k). In contrast, both dexamethasone treatment and PLX5622-mediated depletion markedly reduced the accumulation of vascular CD43⁺ cells compared with lecanemab alone (Fig. 6l, m). Because PLX5622 did not substantially alter circulating immune cell populations, this effect supports a role for brain-resident myeloid cells in the recruitment of peripheral immune cells. In contrast, the dexamethasone effect may reflect depletion of circulating monocytes rather than direct suppression of leukocyte adhesion.

**Figure 6.**
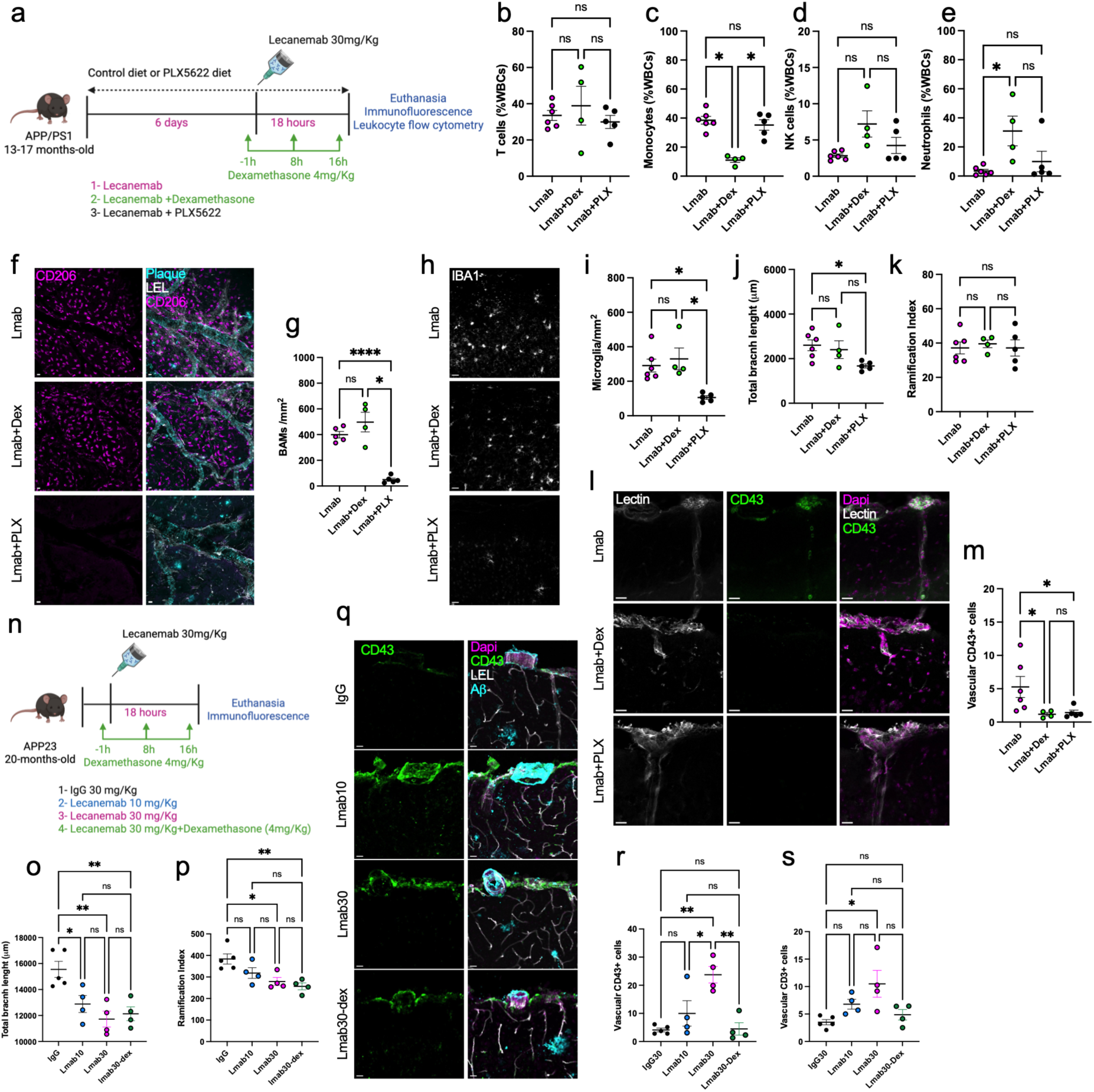
Depletion of brain-resident immune cells and dexamethasone administration attenuate peripheral immune cell recruitment following anti-Aβ immunotherapy. **(a)** Experimental design. APP/PS1 mice (13–17 months old) were maintained on either a control diet or a PLX5622-containing diet for 6 days before lecanemab administration (30 mg/kg). A subset of mice received dexamethasone (4 mg/kg) at −1, 8, and 16 h relative to lecanemab administration. Mice were euthanized 18 h after treatment for immunofluorescence and flow cytometric analysis of leukocytes. **(b–e)** Flow cytometric quantification of circulating T cells (b), monocytes (c), NK cells (d), and neutrophils (e). Dexamethasone significantly reduced circulating monocytes and increased neutrophils, whereas PLX5622 did not significantly alter circulating immune cell populations. Groups were compared using a Kruskal–Wallis test (n = 4-6 per group; overall P = 0.57, 0.007, 0.056, and 0.029, respectively). **(f)** Representative confocal images of CD206+ border-associated macrophages (BAMs). **(g)** Quantification of BAM density demonstrating efficient depletion following PLX5622 treatment, while dexamethasone did not affect BAM density (n = 4-6 per group; Brown-Forsythe ANOVA test, P = 0.005). **(h)** Representative IBA1 immunostaining. PLX5622 markedly reduced the number of microglia **(i)** and total branch length **(j)**, whereas the ramification index was unchanged among the remaining microglia **(k)**. Dexamethasone did not significantly alter microglial morphology or density. Groups were compared using a Kruskal–Wallis test (n = 4-6 per group; P = 0.0015 (i), 0.049 (j), and 0.8 (k)). **(l)** Representative images of vascular CD43+ immune cells in leptomeningeal vessels. **(m)** Quantification of vascular CD43⁺ cells. Cells were counted for each vessel, and a single mean value was calculated for each mouse. Both dexamethasone treatment and PLX5622-mediated depletion significantly reduced lecanemab-induced vascular immune cell adhesion (n = 4-6 per group; Kruskal–Wallis test, P = 0.006). **(n)** Experimental design; APP23 mice (20 months old) received a single injection of IgG (30 mg/kg), lecanemab (10 mg/kg), lecanemab (30 mg/kg), or lecanemab (30 mg/kg) plus dexamethasone (4 mg/kg). Mice were euthanized 18 h later for immunofluorescence analysis. **(o–p)** Quantification of microglial morphology in APP23 mice showing reduced total branch length (o) and ramification index (p) following lecanemab treatment, with no significant changes due to dexamethasone administration (n = 4-5 per group; One-way ANOVA (P = 0.002 and 0.005, respectively). **(q)** Representative images for vascular CD43^+^ cells in APP23 mice. Quantification of vascular CD43^+^ **(r)** and CD3^+^ **(s)** immune cells demonstrating increased vascular immune cell recruitment after 30 mg/Kg lecanemab (n = 4-5 per group; One-way ANOVA, P = 0.0007 and 0.013, respectively). One-way ANOVA was followed by Holm–Šídák’s multiple-comparisons test, Kruskal–Wallis by Dunn’s test, and Brown–Forsythe and Welch ANOVA by Dunnett’s T3 multiple-comparisons test. *P < 0.05, **P < 0.01, ***P < 0.001, ****P < 0.0001; ns, not significant. Scale bars, 20 µm. Schematics in (a) and (n) were created with BioRender.com.

To determine whether these findings extend to a model with robust CAA, aged APP23 mice (20 months old) received IgG, low-dose lecanemab (10 mg/kg), high-dose lecanemab (30 mg/kg), or high-dose lecanemab plus dexamethasone and were analyzed 18 h later (Fig. 6n). High-dose lecanemab induced pronounced microglial activation, characterized by reduced total branch length and decreased ramification index, and dexamethasone did not prevent these morphological changes (Fig. 6o, p), suggesting that microglia remain responsive to antibody-bound amyloid despite systemic immunosuppression. In contrast, dexamethasone significantly reduced the number of adherent vascular CD43⁺ cells following high-dose lecanemab treatment (Fig. 6q-s). Together, these findings demonstrate that brain-resident myeloid cells are critical mediators of peripheral immune cell recruitment following anti-Aβ immunotherapy, whereas acute microglial activation is not affected by dexamethasone treatment.

## Discussion

Anti-Aβ immunotherapies have transformed the treatment landscape for Alzheimer’s disease, yet the biological responses triggered immediately following antibody engagement of amyloid remain poorly understood. While substantial effort has focused on the mechanisms of amyloid clearance, considerably less is known about the earliest neuroimmune responses elicited by treatment. Here, using longitudinal in vivo two-photon imaging and complementary immunohistological and flow cytometric analyses, we demonstrate that anti-Aβ immunotherapy rapidly induces coordinated activation of both resident and peripheral immune cells. Within 24 hours of treatment, aducanumab and lecanemab triggered extensive microglial activation around amyloid plaques, accompanied by endothelial activation and recruitment of monocytes, neutrophils, and T cells to leptomeningeal vessels.

Microglial activation has long been considered a central mechanism underlying antibody-mediated amyloid clearance^2,13^. Recent studies have further demonstrated that lecanemab requires Fc-mediated microglial activation to effectively clear amyloid plaque^13^. Consistent with this model, we observed that anti-Aβ immunotherapy induced rapid morphological activation of plaque-associated microglia and suppressed plaque growth. Microglial responses emerge within 24–48 hours of antibody administration, indicating that microglial engagement is among the earliest biological responses to therapeutic amyloid binding. Depletion of brain-resident myeloid cells attenuated recruitment of peripheral immune cells to leptomeningeal vessels, while dexamethasone decreased the number of adherent peripheral immune cells without preventing microglial activation. Together, these findings support a model in which resident CNS myeloid-cell activation following anti-Aβ immunotherapy suppresses amyloid plaque growth and contributes to signals that promote leukocyte adhesion and recruitment at leptomeningeal vessels.

Although our depletion strategy does not distinguish microglia from BAMs, several observations suggest that BAMs may be particularly important in initiating this response. Peripheral immune-cell recruitment occurred predominantly within leptomeningeal vessels, where BAMs are positioned at the vascular interface^6^, and BAMs were frequently observed in close proximity to adherent peripheral immune cells. Moreover, peripheral immune-cell recruitment was evident before the more pronounced changes in microglial activation. Together, these findings raise the possibility that BAMs provide an early link between antibody engagement of vascular amyloid and peripheral leukocyte recruitment, while microglial activation may contribute to subsequent parenchymal responses and amyloid clearance.

Three observations suggest that the acute vascular immune response identified here represents an early event in the pathogenesis of ARIA: 1) Immune cell recruitment occurred predominantly within leptomeningeal and cortical vessels, the vascular compartments most frequently affected in ARIA^1^. 2) The higher lecanemab dose in APP23 produced a larger peripheral immune response, paralleling the established dose-dependent risk of ARIA observed clinically even after a single antibody dose^14^. 3) Finally, the recruited populations included both monocytes and T cells, consistent with recent multiomic profiling of ARIA-positive patients demonstrating expansion of cytotoxic CD8+ T cells alongside enhanced CD14+ and CD16+ monocyte signaling through antigen presentation, adhesion, and chemokine pathways^5^.

Our brain-resident myeloid cell depletion studies suggest that activation of resident CNS myeloid cells contributes to peripheral immune recruitment, providing a framework for interpreting the sequence of events following anti-Aβ immunotherapy that may lead to ARIA. Building on the model proposed by Taylor et al. ^7^, one plausible mechanism is that increasing antibody concentrations increases CAA-bound antibody in cerebral arteries and soluble or cleared Aβ–antibody complexes in cerebral veins, thereby activating brain-resident immune cells, particularly BAMs. This, in turn, may promote endothelial activation and upregulation of adhesion molecules such as ICAM1, thereby facilitating the recruitment and retention of peripheral immune cells within the cerebrovascular compartment. Recent studies have identified complement activation following antibody binding to cerebrovascular amyloid as one of the earliest events following anti-Aβ immunotherapy^15^. Whether complement activation precedes the cerebrovascular immune response described here or is amplified by recruited peripheral immune cells remains unknown. Defining the temporal and mechanistic relationships among resident CNS myeloid cell activation, complement activation, and peripheral immune cell recruitment will be critical for understanding the initiation of ARIA.

APOE4 is the strongest genetic risk factor for ARIA, although the mechanisms underlying this increased susceptibility remain poorly understood^1^. We show that anti-Aβ immunotherapy increases the fraction of nonclassical monocytes in the peripheral circulation and recruits T cells to leptomeningeal vessels. Recent large-scale proteomic profiling identified an APOE ε4-associated signature enriched for immune and inflammatory pathways across multiple neurodegenerative diseases^16^. These proteomic changes mapped predominantly to nonclassical and intermediate monocytes, memory CD8⁺ T cells, regulatory T cells, and NK/γδ T-cell populations, suggesting a broad immune activation phenotype associated with APOE ε4^16^. Taken together, this raises the possibility that increased ARIA risk in APOE4 carriers may, in part, be explained by a more pronounced pro-inflammatory response involving nonclassical monocytes and CD8⁺ T cells following anti-amyloid immunotherapy.

An intriguing finding from both the mouse and human datasets was the emergence of proliferative myeloid populations following anti-Aβ immunotherapy. In our mouse studies, lecanemab treatment increased the number of vascular-associated IBA1^+^ Ki67^+^ cells. These cells may represent recently recruited bone marrow-derived monocytes adhering to activated vessels. Alternatively, they may correspond to transitional monocytes (TR-Mo), a recently described proliferative intermediate state that emerges during monocyte-to-macrophage differentiation^2,12^. TR-Mo are characterized by a transient proliferative program marked by expression of cell-cycle genes, including MKI67, CDK1, and TOP2A, followed by upregulation of genes involved in cell adhesion, antigen presentation, and tissue engraftment ^12^. This developmental trajectory has been proposed to allow a limited number of recruited monocytes to expand locally before differentiating into tissue-resident macrophages^12^. Consistent with this possibility of emergence of TR-Mo, single-cell transcriptomic analysis of postmortem brain tissue from a patient who died following lecanemab-associated CAA-related inflammation^2,4^ revealed higher TR-Mo signature scores within monocytes compared with untreated AD controls, together with increased expression of proliferation-associated genes. Pathway enrichment and predicted ligand–receptor interactions suggest that monocytes with this signature may participate in antigen presentation and communication with other immune cells. Together with reports of expanded cytotoxic CD8⁺ T cells in patients with ARIA^5^, it is tempting to speculate that TR-Mo may contribute to local antigen presentation and downstream T cell activation.

Our findings identify a rapid peripheral immune response to anti-Aβ immunotherapy that involves multiple immune cell populations and raise important questions about their contributions to vascular injury. Although APP/PS1 and APP23 mice provide complementary models of amyloid pathology and cerebral amyloid angiopathy, differences in Aβ production and accumulation between these models and human Alzheimer’s disease may influence the kinetics and magnitude of the immune response. Moreover, the murine IgG2a surrogates used here preserve activating FcγR engagement but differ from the clinical human IgG1 antibodies in properties that may affect Fc-mediated responses. Extending these findings to patients by analyzing peripheral blood and cerebrospinal fluid collected before and after antibody administration, together with available postmortem tissue from patients who develop ARIA, will be important for defining the specific immune cell populations that contribute to ARIA-associated vascular injury.

Collectively, these findings support a revised model in which anti-Aβ antibodies rapidly initiate a coordinated cerebrovascular immune response involving resident CNS myeloid cells, endothelial activation, and recruitment of peripheral innate and adaptive immune cells. While these responses may facilitate amyloid clearance, they may also represent the earliest biological events that predispose susceptible patients to ARIA. Limiting this acute peripheral inflammatory response while maintaining microglial activation may therefore preserve therapeutic plaque-clearing capacity and reduce treatment-associated vascular complications.

## Methods

### Animals

#### Mice

APPswe/PS1dE9 transgenic mice (APP/PS1; Stock No. 034829, The Jackson Laboratory; C57BL/6;C3H background) were crossed with Tmem119-TdTomato-CreERT2 transgenic mice (gift from the El Khoury laboratory) to enable specific visualization of resident microglia. Two generations of breeding were used to generate hemizygous APP/PS1 mice homozygous for the Tmem119-tdTomato reporter. TMEM119 expression is restricted to resident microglia. APP23 mice were obtained from The Jackson Laboratory (Stock No. 030504) and aged to 20 months prior to experimentation. Both males and females were used. Mice of the same sex were group-housed (up to four per cage) prior to cranial window surgery and singly housed thereafter. Animals were maintained under a 12 h light/dark cycle with ad libitum access to food and water in a pathogen-free facility. All procedures were approved by the MGH Institutional Animal Care and Use Committee.

#### Validation of Tmem119-tdTomato reporter specificity

To validate reporter specificity, Tmem119-tdTomato mice were perfused with PBS followed by 4% paraformaldehyde and brains were collected and post-fixed in 4% paraformaldehyde for 4 h. After antigen retrieval in citrate buffer, coronal brain sections were stained with IBA1 (Synaptic systems, catalog # 234009) and RFP (Rockland, catalog # 600-401-379) and imaged by confocal microscopy. Co-localization of tdTomato fluorescence with IBA1 immunoreactivity was assessed to confirm selective labeling of resident microglia by the Tmem119-tdTomato reporter.

### In vivo two-photon imaging

#### Cranial window surgery

Cranial window surgeries with metal head plates for stable imaging were performed as described previously^17^. Briefly, mice were anesthetized with isoflurane (5% induction, 1.5% maintenance) and placed on a heated stereotaxic frame. A 4 mm craniotomy was performed over the somatosensory cortex and sealed with a sterile 5 mm glass coverslip using dental cement and cyanoacrylate adhesive. A custom metal head plate (Narishige, CP-2) was affixed using C&B Metabond (Parkell) to stabilize the head during imaging. Postoperative care included analgesia with meloxicam and temperature support for three days.

#### In vivo two-photon imaging

In vivo imaging was performed using a Bruker Ultima 2P Plus microscope equipped with galvanometer and resonant scanners, three GaAsP detectors, and a 25× 1.05 NA water-immersion objective (Olympus) coupled to a piezoelectric z-drive. Mice were anesthetized with isoflurane (1.5% maintenance) with the head plate secured in a frame (Narishige, MAG-1). Two-photon excitation was achieved using a tunable Ti:Sapphire laser (800–1050 nm). Emission was collected in blue (400–460 nm), green (480–560 nm), and red (575–630 nm) channels. Laser power was adjusted to avoid saturation, while detector settings were kept constant for each mouse across imaging sessions. To visualize circulating and adherent leukocytes, 50 μL of sodium azide-free Lycopersicon esculentum lectin (LEL) was administered by retro-orbital injection immediately before baseline imaging and once daily thereafter. tdTomato-positive microglia were visualized using 1050 nm excitation, whereas methoxy-X04-labeled amyloid plaques were imaged using 800 nm excitation. For vascular imaging experiments, LEL-labeled leukocytes were visualized using 900 nm excitation. Z-stacks were acquired at approximately one frame per second. Rolling cells were defined as lectin-positive cells moving along the vessel wall between consecutive frames, whereas adherent cells remained stationary throughout the entire z-stack acquisition. Representative time-series movies of rolling cells were acquired in a single imaging plane at 0.8 seconds per frame. The number of vascular adhering and rolling cells was counted per field of view and averaged per mouse.

### Experimental design and treatment

#### Experimental cohorts

Independent cohorts of mice were used for (i) weekly longitudinal aducanumab imaging experiments, including subsequent treatment-switching studies in the IgG-treated animals, (ii) daily longitudinal aducanumab and lecanemab imaging experiments, including subsequent lecanemab treatment-switching studies in the IgG-treated animals, (iii) acute postmortem analyses following lecanemab treatment, (iv) APP/PS1 studies involving dexamethasone administration or microglial depletion with PLX5622, and (v) APP23 studies examining dose-dependent effects of lecanemab and dexamethasone co-treatment. Aducanumab (mouse IgG2a), the lecanemab surrogate mAb158 (mouse IgG2a), and their respective matched isotype controls are commercially available. mAb158 (Cat. No. HY-P990110) and its matched mouse IgG2a κ isotype control (Cat. No. HY-P99978) were purchased from MedChemExpress. Antibodies and their respective isotype controls were administered intraperitoneally at doses of 10 or 30 mg/kg.

#### In vivo monitoring of microglia and vasculature after aducanumab administration

Microglia were visualized via tdTomato fluorescence, and amyloid plaques were labeled by intraperitoneal injection of methoxy-X04 (∼5 mg/kg) 24 h before each imaging session. Two longitudinal imaging paradigms were employed. For weekly imaging experiments, 9-12 months old APP/PS1-Tmem119-tdTomato mice received cranial window surgery and head plate implantation. APP/PS1-Tmem119-tdTomato mice were allowed to recover for 4 weeks, then received weekly injections of aducanumab (10 mg/kg, n=4) or IgG control (n=3) and were imaged at baseline and weekly for 3 weeks, with antibody administration immediately following each imaging session. To control for baseline plaque burden and inter-animal variability, a subset of mice initially treated with IgG was subsequently switched to aducanumab. APP/PS1-Tmem119-tdTomato mice underwent baseline imaging, followed by weekly IgG administration and imaging for 3 weeks; the same animals then received weekly aducanumab and were imaged for an additional 3 weeks. For daily imaging experiments, mice received a single injection of aducanumab (10 or 30 mg/kg, n= 3 per group) or matched IgG control (n=2-3) and were imaged at baseline, Day 1, Day 2, Day 5, and Day7 post-treatment. Identical plaque-associated fields of view were re-identified using vascular landmarks and cranial window coordinates, allowing within-animal comparisons of microglial dynamics before and after aducanumab treatment.

For in vivo vascular imaging after aducanumab administration, blood vessels were labeled by retro-orbital injection of fluorescein-dextran (70 kDa, 12.5 mg/mL) and Texas Red-dextran (3 kDa, 12.5 mg/mL) immediately before baseline imaging. The same vascular regions were re-imaged 18 h after antibody administration to quantify leukocyte rolling, adhesion, and vascular immune responses.

#### In vivo monitoring of microglia and vasculature after lecanemab administration

9-12-month-old APP/PS1-Tmem119-tdTomato mice underwent cranial window surgery and head plate implantation. After 4 weeks of recovery, mice were imaged at baseline and at 7, 18, and 48 hours following a single injection of lecanemab (30 mg/kg, n= 3) or IgG (30 mg/kg, n=4). To determine whether the rapid increase in intravascular leukocyte recruitment following lecanemab treatment was attributable to antibody administration rather than baseline vascular characteristics or inter-animal variability, mice initially assigned to the IgG treatment group were followed longitudinally and subsequently switched to lecanemab treatment. APP/PS1-Tmem119-tdTomato mice received IgG and underwent in vivo imaging at 7 h, Day 1, and Day 2 after treatment, followed by an additional imaging session on Day 5. The same animals were then switched to lecanemab treatment on Day 5 and re-imaged at 7 h, Day 6, and Day 7 (corresponding to 7 h, Day 1, and Day 2 after lecanemab administration). The same vascular segments were re-identified across imaging sessions using vascular branching patterns and cranial window landmarks. Rolling and adherent lectin-positive intravascular cells were quantified throughout the experiment as described above, enabling within-animal comparisons before and after initiation of lecanemab treatment. Rolling intravascular cells were defined as lectin-positive cells moving along the vessel wall between consecutive image frames within a z-stack, whereas adherent intravascular cells were defined as lectin-positive cells that remained stationary throughout the entire z-stack acquisition.

#### Lecanemab treatment in APP/PS1 mice for postmortem experiments

A separate cohort of APP/PS1 mice received 30 mg/Kg lecanemab (n=6) or IgG (n=4) control and were euthanized 18 h after treatment for immunofluorescence-based characterization of vascular and parenchymal immune cell populations.

#### PLX5622 and dexamethasone treatment paradigm in APP/PS1 mice

APP/PS1 mice (13–17 months old) were randomly assigned to one of three treatment groups: 30 mg/Kg lecanemab alone (n=6), 30 mg/Kg lecanemab plus dexamethasone (n=5), or 30 mg/Kg lecanemab following microglial depletion with PLX5622 (n=5). For microglial and border-associated macrophage depletion, mice received PLX5622-formulated chow (1200 ppm; Research Diets Inc.) for 6 days prior to antibody administration and throughout the experiment. Control animals received standard chow. Lecanemab was administered as a single intraperitoneal injection (30 mg/kg), and mice were euthanized ∼18 hours later for tissue collection. For corticosteroid treatment, dexamethasone (4 mg/kg, i.p.) was administered 1 hour before lecanemab injection and repeated at 8 and 16 hours following lecanemab administration. Mice were euthanized 18 to 22 hours after lecanemab treatment. Brains and blood were collected for immunofluorescence analysis and for counting circulating leukocytes by flow cytometry.

#### Dexamethasone treatment paradigm in APP23 mice

APP23 mice (20 months old) were randomly assigned to receive a single intraperitoneal injection of IgG control (30 mg/kg, n=5), lecanemab (10 mg/kg, n=4), lecanemab (30 mg/kg, n=4), or lecanemab (30 mg/kg) combined with dexamethasone treatment (n=4). Dexamethasone (4 mg/kg, i.p.) was administered 1 hour before lecanemab injection and subsequently at 8 and 16 hours after treatment. Animals were euthanized 18 to 22 hours after antibody administration, and brains were processed for immunofluorescence staining.

### Tissue Collection and Histology

#### Postmortem tissue processing

Mice were euthanized with carbon dioxide and immediately perfused intracardially with ice-cold phosphate-buffered saline (PBS). Brains were removed and hemisected. One hemisphere was post-fixed in 4% PFA for 4 hours at 4°C, cryoprotected in 15% and 30% sucrose, embedded, and sectioned coronally at 40 μm using a vibratome for immunofluorescence staining and stored in cryoprotectant at −20 °C. The contralateral hemisphere was processed for whole-mount analysis of leptomeningeal vessels and border-associated macrophages. For APP23 mice, LEL was administered by retro-orbital injection immediately before euthanasia to label the vasculature.

#### Immunofluorescence staining

Mounted or free-floating sections were washed in Tris-buffered saline (TBS) and blocked in TBS containing 10% normal goat or donkey serum. For CD43, CD3, Ly6G, anti-Aβ and ICAM1 staining, sections were incubated with primary antibodies for 1 h at room temperature without antigen retrieval. For IBA1 and Ki67 staining, antigen retrieval was performed in citrate buffer before blocking and primary antibody incubation. Alexa Fluor-conjugated secondary antibodies were used for visualization. To assess antibody localization, sections were stained with goat anti-mouse IgG2 antibodies to detect lecanemab. Arteries were identified by α-smooth muscle actin (αSMA) immunoreactivity and vessel morphology. CAA-affected vessels were identified by vascular amyloid deposition visualized by methoxy-X04 or anti-Aβ staining.

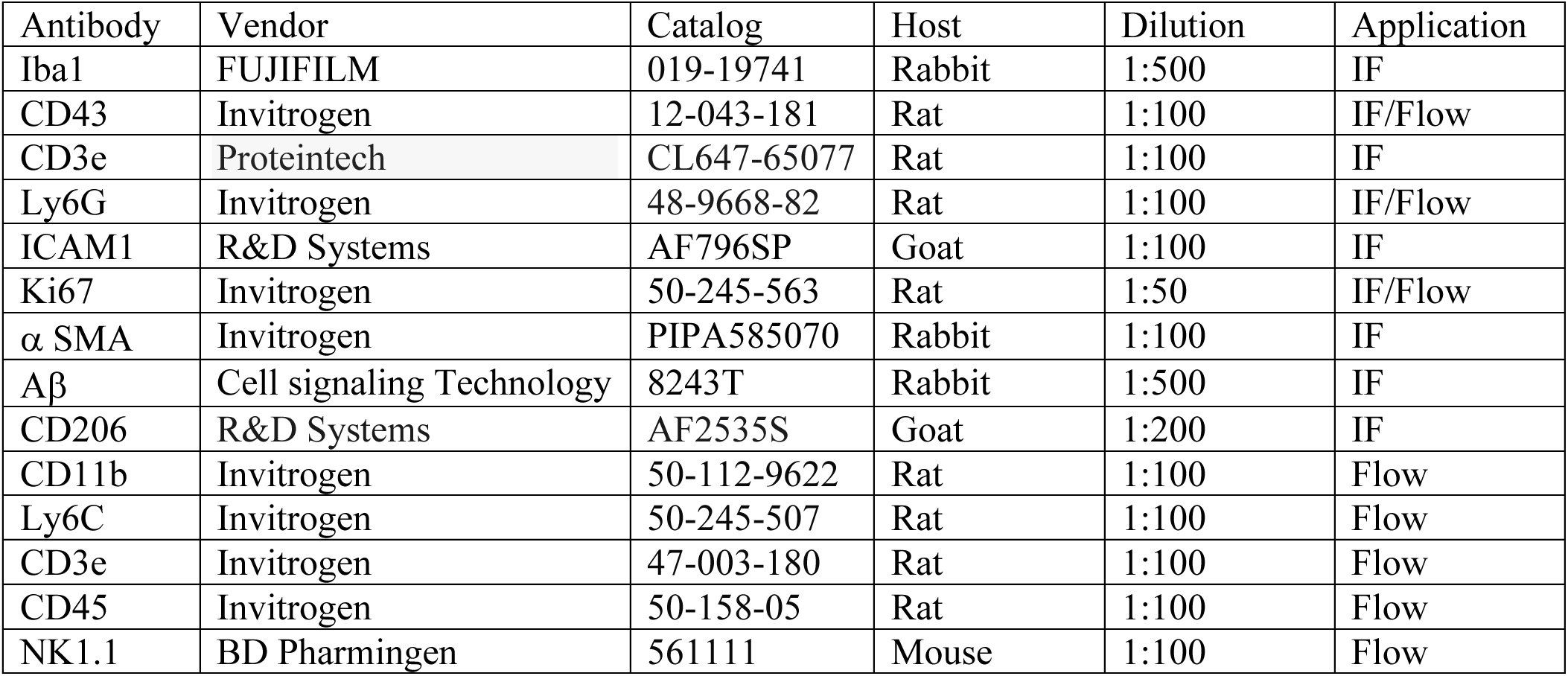

| Antibody | Vendor | Catalog | Host | Dilution | Application |
| --- | --- | --- | --- | --- | --- |
| Iba1 | FUJIFILM | 019-19741 | Rabbit | 1:500 | IF |
| CD43 | Invitrogen | 12-043-181 | Rat | 1:100 | IF/Flow |
| CD3e | Proteintech | CL647-65077 | Rat | 1:100 | IF |
| Ly6G | Invitrogen | 48-9668-82 | Rat | 1:100 | IF/Flow |
| ICAM1 | R&D Systems | AF796SP | Goat | 1:100 | IF |
| Ki67 | Invitrogen | 50-245-563 | Rat | 1:50 | IF/Flow |
| α SMA | Invitrogen | PIPA585070 | Rabbit | 1:100 | IF |
| Aβ | Cell signaling Technology | 8243T | Rabbit | 1:500 | IF |
| CD206 | R&D Systems | AF2535S | Goat | 1:200 | IF |
| CD11b | Invitrogen | 50-112-9622 | Rat | 1:100 | Flow |
| Ly6C | Invitrogen | 50-245-507 | Rat | 1:100 | Flow |
| CD3e | Invitrogen | 47-003-180 | Rat | 1:100 | Flow |
| CD45 | Invitrogen | 50-158-05 | Rat | 1:100 | Flow |
| NK1.1 | BD Pharmingen | 561111 | Mouse | 1:100 | Flow |

For whole-mount preparations, hemispheres were fixed in 4% PFA for 2–4 h, blocked and permeabilized in TBS containing 10% bovine serum albumin (BSA) and 0.1% Triton X-100, and incubated with primary antibodies against Border-associated macrophages (CD206) and Aβ for 2–3 days at 4°C. Tissues were then washed and incubated with the appropriate secondary antibodies for 24 h at 4°C. LEL was injected retro-orbitally immediately before euthanasia to label blood vessels. Images were acquired using an Olympus FV3000 confocal microscope equipped with 20× and 40× objectives. Identical acquisition settings were maintained across treatment groups within each experiment.

#### Flow Cytometry

Peripheral blood was collected into K3-EDTA tubes (Sarstedt) and processed on the day of collection. Red blood cells were lysed using RBC lysis buffer (Thermo Fisher) according to the manufacturer’s instructions. Following lysis, cells were washed with PBS containing 2% fetal bovine serum (FBS). Cells were stained with LIVE/DEAD™ Fixable Aqua Viability Dye (Thermo Fisher Scientific) to determine the live cells. Cells were subsequently incubated with pre-titrated fluorophore-conjugated antibodies directed against leukocyte lineage markers and monocyte subset markers for 30 min at 4°C protected from light. Following surface staining, cells were fixed and permeabilized using the BD Cytofix/Cytoperm™ Plus kit (BD Biosciences) and stained intracellularly with anti-Ki67 antibody. Unstained and fluorescence-minus-one controls were used to define gating thresholds. Single-color compensation controls were prepared using BD CompBeads or BD ArC™ compensation beads. Data acquisitions were performed on a SORP 5-Laser BD LSRFortessa X-20 (Flow and Mass Cytometry Core Facility, Massachusetts General Hospital) using compensation matrices generated on the day of acquisition. Data were analyzed using FlowJo software (v10.10.1, BD Biosciences). Leukocyte populations were identified by sequential gating on FSC (Forward Scatter) / SSC (Side Scatter), singlets, live cells, and white blood cells (WBCs). Neutrophils, T cells, NK cells, and monocytes were identified using established marker combinations. Monocyte subsets were further classified according to Ly6C expression as Ly6C^high and Ly6C^low populations. Cellular proliferation was assessed by quantifying the percentage of Ki67-positive cells within total monocytes and Ly6C-defined monocyte subsets. Population frequencies are reported as a proportion of total white blood cells unless otherwise indicated.

### Analysis

#### Microglial dynamics and morphological quantification in vivo

Microglial spatial dynamics were quantified using 612 × 612 µm or 306 × 306 µm fields of view, registered across imaging sessions. For each imaging interval, microglia that disappeared from their prior location (“lost”) and microglia that newly appeared (“appeared”) were manually identified by a blinded observer. The percentage of dynamic microglia was calculated as the average of lost and appeared cells relative to the total microglial population within the field of view. Plaque-associated microglial dynamics were assessed by quantifying dynamic microglia located within 50 µm of the perimeter of methoxy-X04-positive amyloid plaques.

For morphological classification, microglia were manually classified by a blinded observer using images acquired at 4× magnification from the same field of view over time (153 × 153 µm). Microglia were classified as ramified (small soma with thin, highly branched processes), reactive (enlarged soma with thicker and less-branched processes), rod-shaped (elongated soma with polarized processes), or amoeboid (rounded enlarged soma with markedly reduced or absent processes) based on established morphological criteria^10^. The proportion of non-ramified microglia (rod-shaped, reactive and amoeboid) was calculated for each imaging session and compared relative to baseline within each treatment group.

#### Plaque size quantification in vivo

Longitudinal amyloid plaque measurements were performed using methoxy-X04-labeled plaques imaged by in vivo two-photon microscopy. For each plaque, z-stacks were matched across imaging sessions using identical start and end piezo positions to ensure imaging of the same volume over time. Average-intensity projections were generated from matched z-stacks (40–80 μm depth) and used to measure plaque area. Plaque boundaries were manually delineated in Fiji (ImageJ) by a blinded observer, and plaque area was quantified at each time point. Plaque size was normalized to baseline values for each plaque to assess longitudinal changes following treatment. Analyses were performed in APP/PS1 mice receiving weekly injections of aducanumab (10 mg/kg) or an IgG control for 3 weeks, followed by longitudinal imaging over the same period (n = 3 mice per treatment group).

#### Histological quantification of vascular immune cells

Quantification was performed by blinded observers using Fiji (ImageJ). For vascular immune cell analyses, leptomeningeal and cortical vessels were identified based on vascular morphology and lectin labeling. CD43+, CD3+, and Ly6G+ cells were manually quantified per vessel. To be considered positive, cells were required to exhibit circumferential or membrane-associated immunoreactivity surrounding a DAPI-positive nucleus. Structures exhibiting only nuclear immunoreactivity without corresponding membrane staining were considered nonspecific and excluded from analysis. Similarly, immunopositive structures lacking an associated DAPI-positive nucleus were not counted. Ki67⁺ cells were identified by nuclear Ki67 immunoreactivity overlapping with DAPI. IBA1⁺Ki67⁺ cells were identified by a Ki67-positive nucleus within an IBA1-positive cell body. Cell counts from individual vessels were averaged to obtain a single value per mouse, which served as the biological replicate for statistical analyses of adherent cells. For vascular ICAM1 quantification, vascular regions of interest were manually defined, and mean fluorescence intensity per leptomeningeal vessel was measured using Fiji, then averaged per mouse.

Microglia morphology: Microglial morphology was quantified from postmortem immunofluorescence images using Fiji (ImageJ) by a blinded observer. Individual microglial somas were manually outlined to generate regions of interest (ROIs), and soma area was calculated from the enclosed area. Images were thresholded and skeletonized using Fiji, and total process length was measured from the resulting skeletonized images. The ramification index (RI) was calculated as total process length per image divided by soma area, with lower values indicating a more activated morphology characterized by process retraction and soma enlargement.

#### Public post-mortem brain transcriptomic analysis

We analyzed publicly available single-cell transcriptomic data from post-mortem brain tissue obtained from one lecanemab-treated and three untreated Alzheimer’s disease patients^2^. The Cell Ranger count matrices served as input to downstream analysis in Seurat (version 5.1.0)^18^, initially filtering cells with unique molecular identifiers over 5,000 or less than 500, as well as outliers with more than 5% mitochondrial counts. The feature expression measurements for each cell were normalized by the total expression, multiplied by the scale factor (10,000), and log-transformed. We calculated the 3,000 highly variable genes to be used as integration anchors between the samples, followed by linear transformation (scaling) regressing out heterogeneity associated with mitochondrial contamination and UMI counts. Linear dimensionality reduction was performed by PCA, followed by non-linear UMAP embedding using the first 40 principal components; the same parameters were used for clustering with Seurat’s FindNeighbors and FindClusters functions (0.9 resolution). Differential gene expression comparison between groups was performed using the nonparametric Wilcoxon rank-sum test. Genes with fold change > 1.5 (absolute value), adjusted P-value (Bonferroni correction) < 0.05, and detected in a minimum fraction of 20% of cells in either of the two populations were considered differentially expressed. We generated a gene module score for TR-Mo signature (MKI67, TOP2A, CDK1, PCLAF, UBE2C, BIRC5, HIST1H2AE, STMN1, HMGB2, TUBB5, H2AC8) using the Seurat’s function AddModuleScore^19^

#### Single-cell RNA sequencing of mouse blood monocytes

Fresh peripheral blood was collected from APP/PS1 mice 16–18 h after administration of lecanemab or IgG control (n = 3 mice per group). Monocytes were isolated on the day of collection by negative selection using a mouse monocyte isolation kit (STEMCELL Technologies) and submitted to the Center for Cellular Profiling (CCP) for single-cell RNA sequencing using the 10x Genomics platform. Samples from four mice were multiplexed per chip using sample-specific barcodes, targeting 5,000 cells per mouse. Sequencing data were processed using Cell Ranger (version 9.0.1), and sample barcodes were used to assign cells to individual mice. UMAP visualizations were generated in Loupe Browser, and classical and non-classical monocyte populations were annotated based on canonical marker expression. Cells assigned to the platelet cluster were excluded from monocyte quantification. Monocyte subset proportions were calculated separately for each mouse relative to the total number of classical and non-classical monocytes.

#### Statistics

All quantifications were performed by blinded observers. Statistical analyses were conducted using GraphPad Prism (Version 11). Data are presented as mean ± SEM. Normality was assessed using the Shapiro–Wilk test. Parametric data were analyzed using paired or unpaired two-sided t-tests, one-way or two-way ANOVA, repeated-measures ANOVA, or mixed-effects models, as appropriate. Non-parametric data were analyzed using Mann–Whitney U or Kruskal–Wallis tests. Where appropriate, multiple comparisons were adjusted using Holm–Šídák, Tukey’s or Dunn’s tests. Statistical details and sample sizes are reported in the figure legends, and P < 0.05 was considered statistically significant.

## Data availability

All data needed to support the conclusions of this article are provided in the paper and Extended Data. All images will be made available upon reasonable request.

## Acknowledgements

Research reported in this publication was supported by the National Institute on Aging of the National Institutes of Health under Award Numbers R01AG054598-06 and K99AG088296, and by the National Institute of Neurological Disorders and Stroke under Award Number K08NS131530. We gratefully acknowledge the donors of the Alzheimer’s Association for their support through grant 23AARF-1028885.

## Author contributions

M.A. and B.J.B. conceived and designed the study. M.A., B.J.B., and M.G.K. secured funding. M.G.K. and B.J.B. supervised the study. M.A., S.Y., W.N., S.A., L.P., A.R., R.G., and Y.L. collected experimental data. M.A., S.A., M.V.S.-M., L.P., K.R.W., K.K., and S.K. analyzed the data. S.I. and P.d.S. performed the human transcriptomic analysis. J.E.K. provided the Tmem119-tdTomato mouse model. M.A. wrote the initial manuscript draft. S.I., R.G., B.J.B., M.G.K., and M.A. revised the manuscript.

---

**Extended Data Figure 1.**
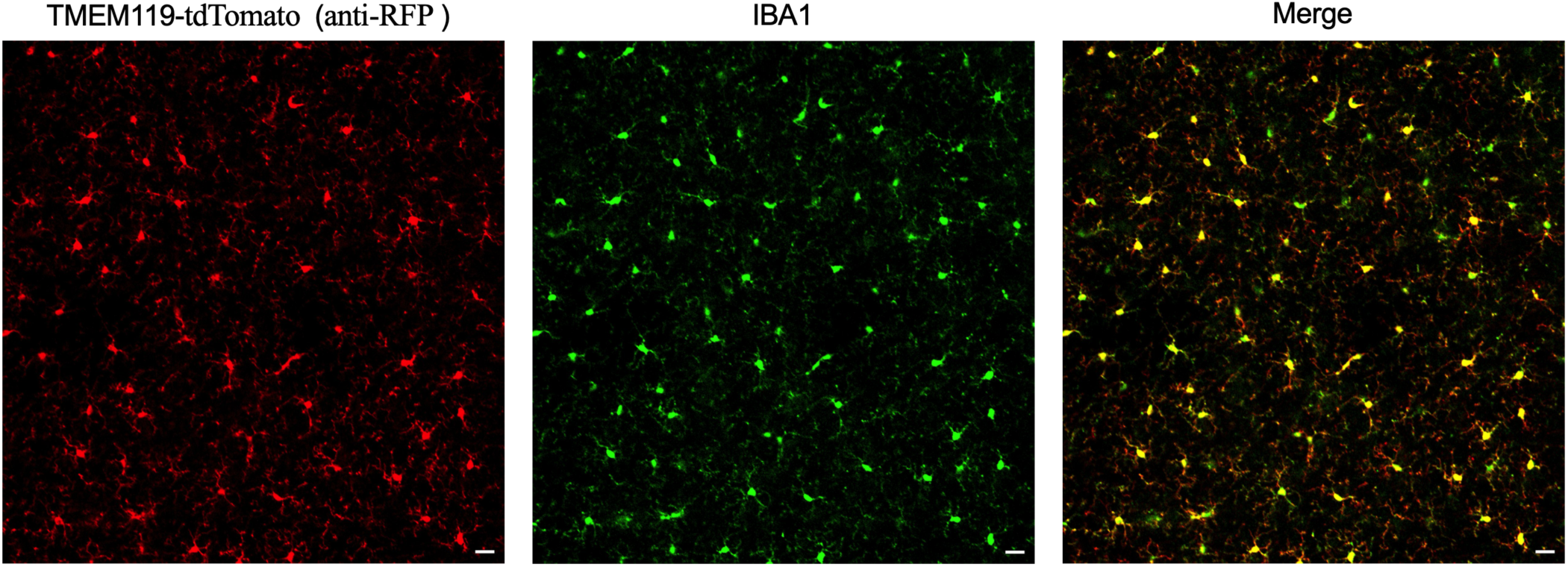
Tmem119-positive microglial cells expressing tdTomato also co-express IBA1. Representative images showing tdTomato (anti-RFP) in Tmem119-expressing microglia, with co-localization of the microglial marker IBA1, confirming the specificity of the Tmem119-tdTomato reporter line. Scale bars, 20 µm.

**Extended Data Figure 2.**
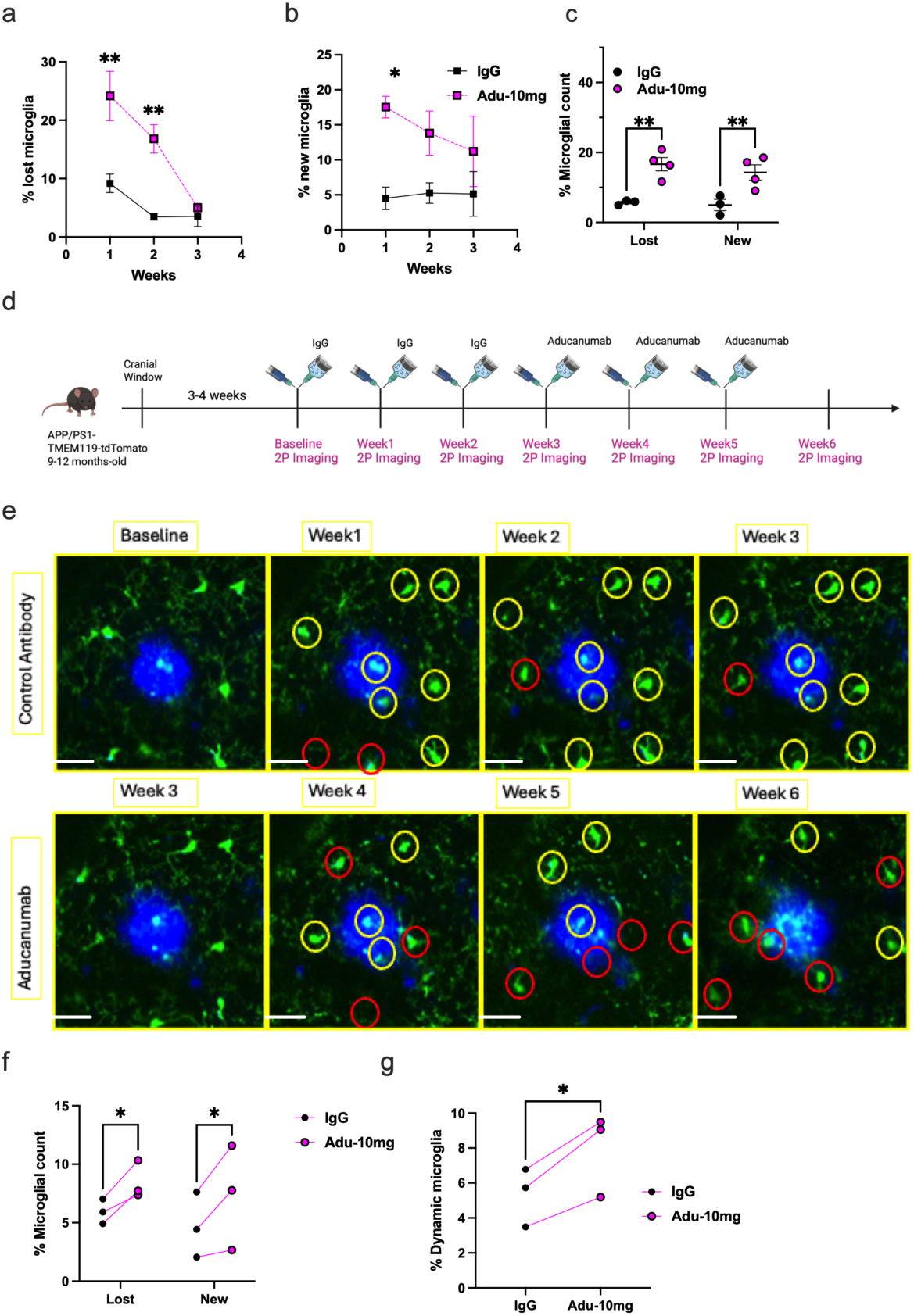
Aducanumab induces increased microglial dynamics within the same plaque-associated field of view. (a, b) Quantification of the percentage of microglia that were lost from their original location (Two-way repeated-measures ANOVA, main treatment effect; P=0.001, n= 3-4 mice per group) or newly appeared within the imaging field (Two-way repeated-measures ANOVA, main treatment effect; P=0.038, n= 3-4 mice per group) over 3 weeks of longitudinal imaging in APP/PS1 mice treated with IgG or aducanumab. (c) Average weekly percentage of lost or newly appearing microglia in APP/PS1 mice receiving IgG or aducanumab (Two-way repeated-measures ANOVA, main treatment effect; P=0.0007, n= 3-4 mice per group). (d) Timeline for longitudinal imaging before and after switching from IgG to aducanumab. Following baseline imaging, APP/PS1-TMEM119-tdTomato mice received weekly IgG for 3 weeks and were then switched to weekly aducanumab for an additional 3 weeks. Two-photon imaging was performed weekly through week 6. These were the same three mice included in the IgG-treated group in Figure 1. (e) Representative images of the same plaque in an IgG-treated mouse that was switched to aducanumab after the third IgG dose (green: microglia, blue: amyloid plaque). Microglia are color-coded based on spatial stability: yellow denotes stable microglia maintained across imaging sessions, while red denotes dynamic microglia that either disappeared from or newly appeared within the imaging field between sessions. (f) Quantification of the percentage of microglia lost or newly appearing during IgG treatment and after switching to aducanumab within the same field of view (Two-way repeated-measures ANOVA, main treatment effect; P=0.016), n= 3 mice per group. (g) Quantification of microglial dynamics across three mice shows low turnover during IgG treatment followed by a marked increase after switching to aducanumab, indicating enhanced microglial dynamics within the same plaque-associated regions (Paired two-sided *t*-test, *P* = 0.032). Two-way ANOVA was followed by Holm–Šídák’s multiple-comparisons test. Scale bars, 20 µm. \**P* < 0.05, \*\**P* < 0.01, \*\*\**P* < 0.001. The schematic in (d) was created with BioRender.com.

**Extended Data Figure 3.**
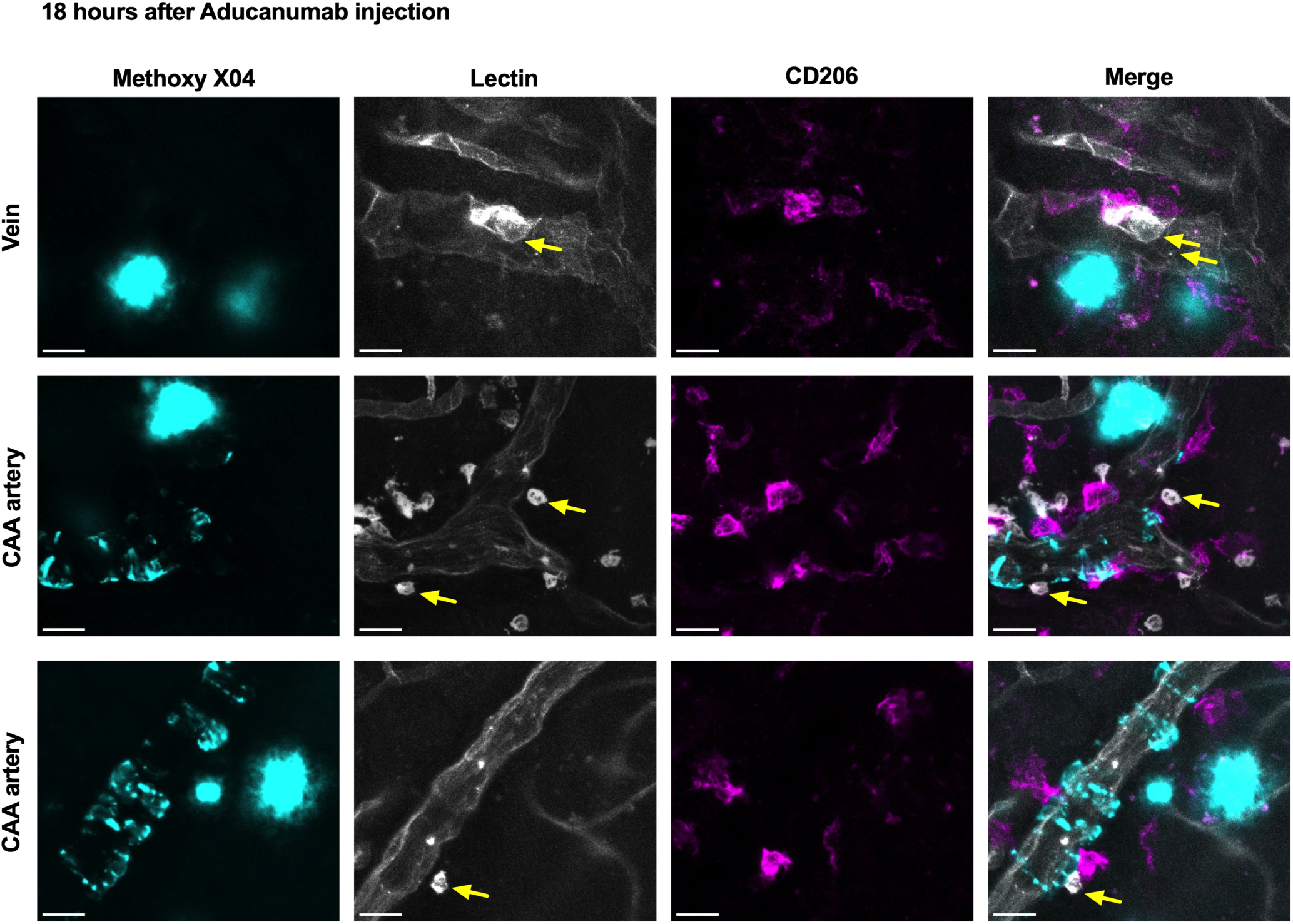
Perivascular macrophages localize near infiltrating lectin-positive peripheral immune cells following aducanumab treatment. Representative ex-vivo confocal images acquired 18 h after aducanumab injection show amyloid plaques labeled with Methoxy-X04 (cyan), vasculature and peripheral immune cells labeled with Lycopersicon esculentum (LE) lectin (white), and border-associated macrophages (BAMs; CD206, magenta). Top row: meningeal vein; middle and bottom rows: CAA-affected arteries. In veins, BAMs are enriched along the vessel wall and are frequently found in close proximity to adhering LE lectin–positive cells. In contrast, in CAA-affected arteries, lectin-positive immune cells can occasionally be observed outside the vessel wall and adjacent to BAMs (yellow arrows). These observations suggest that vascular sites of peripheral immune cell adhesion are associated with BAM recruitment in veins, whereas this process appears to be altered in CAA-affected arteries. Scale bars, 20 µm.

**Extended Data Figure 4.**
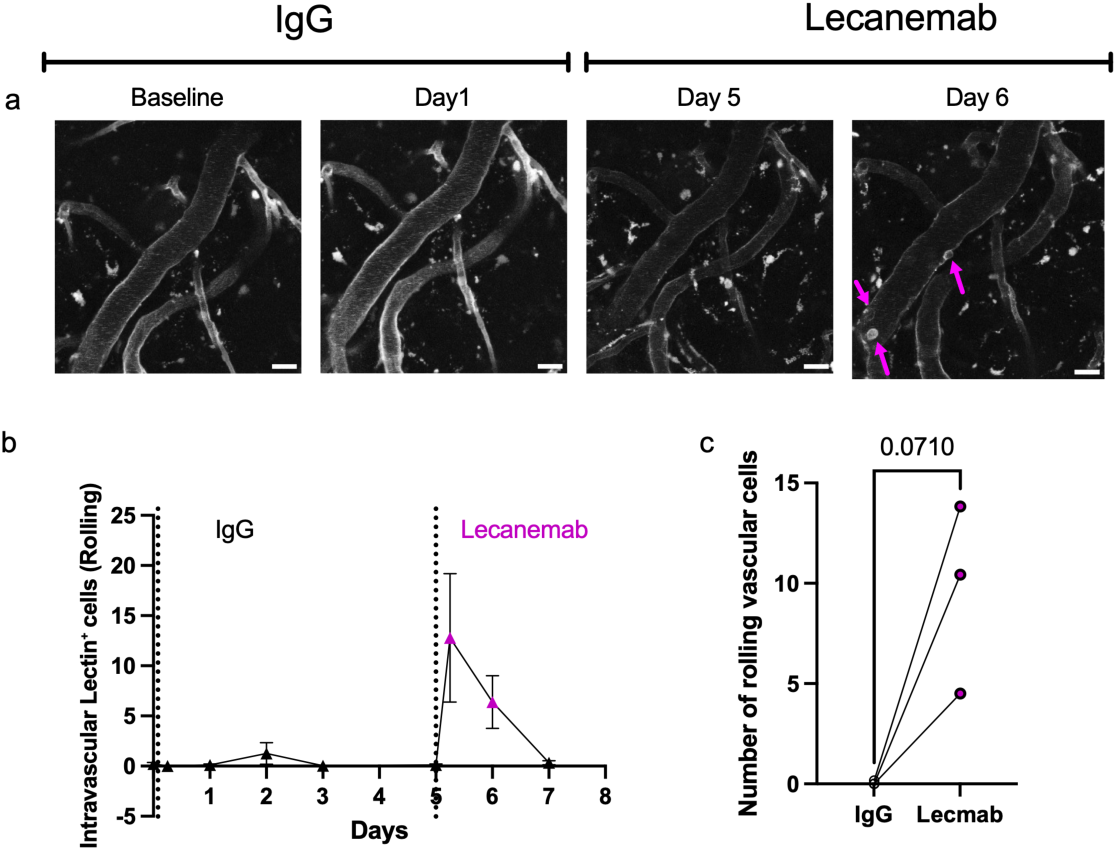
Intravascular rolling and adhesion of leucocytes following lecanemab administration. APP/PS1 mice were longitudinally imaged by in vivo two-photon microscopy before and after switching from IgG to lecanemab treatment. (a) Representative images from the same vessel show intravascular rolling cells after lecanemab administration (magenta arrows). (b)Average number of intravascular lectin+ rolling cells in APP/PS1 mice longitudinally imaged during IgG treatment for 5 days, followed by switching to lecanemab and re-imaging for an additional 2 days. (c) Quantification of the average rolling cells on Day 1 after IgG versus lecanemab treatment in the same mice revealed minimal rolling cells during IgG treatment, followed by a trend toward increased rolling after lecanemab administration within the same vessels (Paired two-sided *t*-test, *P* = 0.07). n = 3 mice. Scale bars, 20 µm.

**Extended Data Fig. 5.**
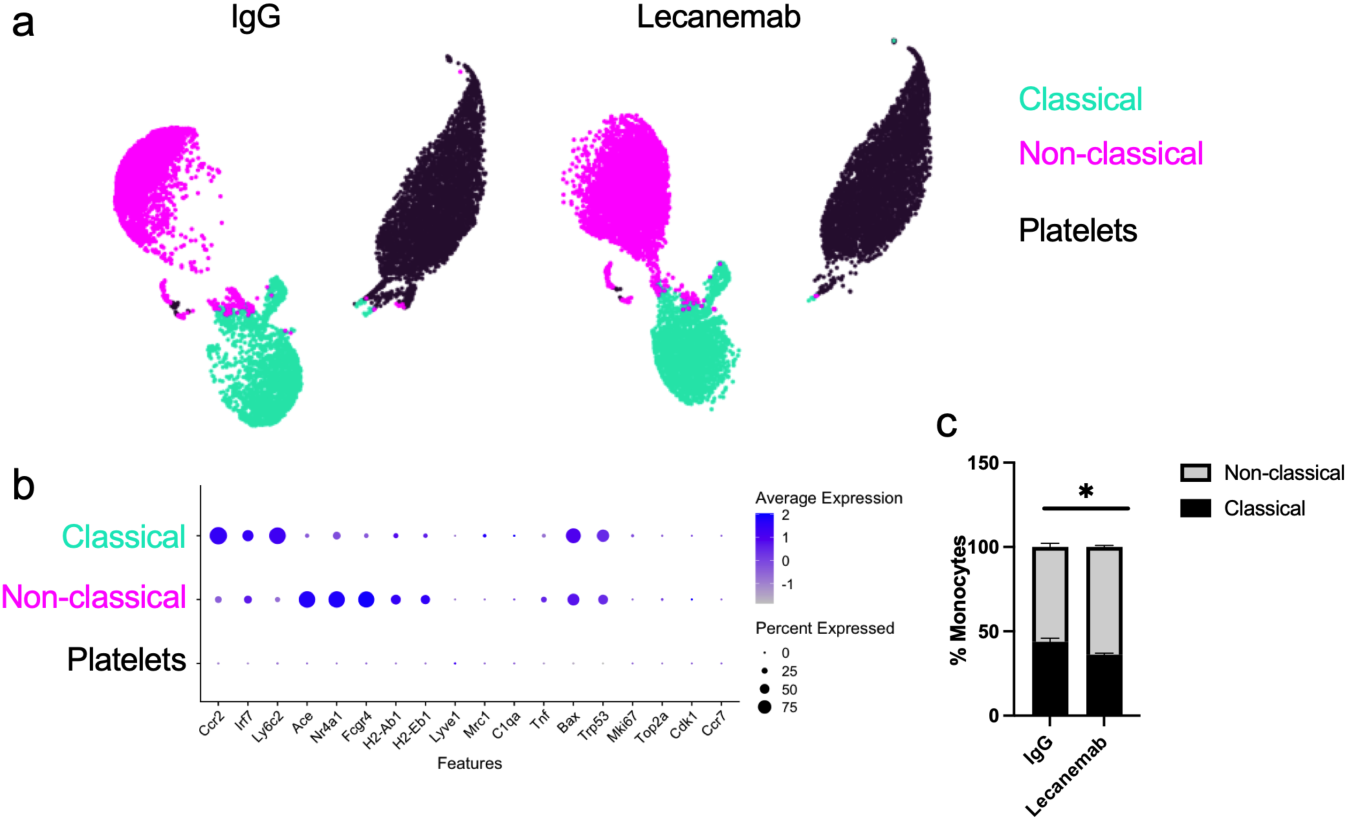
Lecanemab treatment increases the proportion of circulating non-classical monocytes. **(a)** UMAP visualization of negatively isolated blood monocytes collected 16– 18 h after IgG or lecanemab treatment, showing segregation into classical (cyan), non-classical (magenta), and platelet (black) clusters. **(b)** Dot plot showing the expression of canonical marker genes used to identify classical and non-classical monocyte populations. Dot size represents the percentage of cells expressing each gene, and color indicates average expression level. (c) The proportion of non-classical monocytes increased following lecanemab treatment compared with IgG controls (unpaired two-sided t-test, P=0.03, n=3 mice per group).

